# sctrial: Participant-Level Differential Analysis for Longitudinal Single-Cell Experiments

**DOI:** 10.64898/2026.04.02.716219

**Authors:** Priyanka Vasanthakumari, Itzel Valencia, Maryam Ranjpour Aghmiouni, Bryan Magana, Mohamed Omar

**Author notes:** Correspondence to: Mohamed Omar, MD, Assistant Professor, Department of Computational Biomedicine Department of Urology, Cedars-Sinai Medical Center, 700 N. San Vicente Blvd, G-540F West Hollywood, CA 90069, USA.

## Abstract

Longitudinal single-cell RNA sequencing studies in clinical trials and translational cohorts offer a powerful view of treatment response, disease progression, and cellular dynamics, but their hierarchical structure poses a major inferential challenge: thousands of cells are measured within the same participants across repeated time points, whereas the participant, not the cell, is the true unit of biological replication. Conventional cell-level workflows can therefore yield inflated significance and misleading confidence in reported associations. Here, we present *sctrial*, an open source analytical framework for repeated-measures single-cell studies that uses design-specific participant-level estimands, including difference-in-differences for two-group longitudinal comparisons, and small-cluster-aware uncertainty quantification. In simulation benchmarks using a hierarchical gamma-Poisson generative model, sctrial maintained well-calibrated error rates in mixed-signal gene panels where established multi-subject methods showed inflated false positive rates among unaffected genes. We applied *sctrial* to five independent datasets spanning melanoma immunotherapy, COVID-19 severity, BNT162b2 vaccination, AML chemotherapy, and CAR-T therapy. Across these studies, *sctrial* identified immune programs whose direction and magnitude differed across therapeutic and disease contexts, while benchmarking analyses showed that many associations highlighted by conventional cell-level workflows were attenuated or no longer supported when inference was performed at the participant level. These analyses illustrate how participant-aware inference can reduce pseudoreplication-driven signal inflation and provide a more rigorous basis for interpreting longitudinal single-cell data. *sctrial* enables reproducible participant-level analysis of longitudinal single-cell experiments and facilitates more reliable biological interpretation in translational and clinical studies. The software is implemented in Python and compatible the AnnData/scverse ecosystem.

## INTRODUCTION

The advent of single-cell RNA sequencing (scRNA-seq) has transformed the study of cellular heterogeneity by enabling transcriptomic profiling at single-cell resolution across complex tissues, developmental systems, and disease states^1,2^. Increasingly, scRNA-seq is being applied in longitudinal and repeated sampling designs, including clinical trials and translational cohort studies, in which participants are profiled across treatment, disease course, or other clinically meaningful intervals^3^. These designs are common in investigations of cancer immunotherapy^4–6^, vaccination^7–9^, infectious disease^10–14^, and hematologic malignancy^15,16^, where the central objective is to determine how cell states, cell composition, and molecular programs change within participants over time and whether those changes differ across clinically relevant groups.

Despite the growing prevalence of such studies, many single-cell datasets are still analyzed using statistical procedures that do not respect their hierarchical structure. In these data, thousands of cells are measured within the same participant yet cells are frequently treated as independent observations, creating pseudoreplication that overstates the effective sample size^17,18^. Cells from the same individual share genetic background, environmental exposures, clinical context, and technical sources of variation, inducing within-participant correlation that violates the independence assumptions of standard cell-level tests and can inflate statistical significance beyond what the data support^19^. Longitudinal designs introduce an additional inferential challenge^20^. In many studies, the primary question is not whether two groups differ at a single time point, but whether they change differently over time, and analyses that collapse or ignore repeated measurements obscure this structure^21^.

Several statistical strategies have been developed to address aspects of multi-subject single-cell inference^22,23^. Pseudobulk approaches aggregate measurements to the replicate level within a cell population and then apply sample level differential expression methods, thereby aligning inference with the biological unit of replication^24–26^. Mixed-effects models instead represent the hierarchical structure directly, including subject-aware negative-binomial formulations such as NEBULA^27^. These approaches are important and often effective, but they are oriented primarily toward cross sectional group comparisons. Although the underlying linear model frameworks can in principle accommodate longitudinal designs through interaction terms and subject blocking, this requires manual specification of design matrices, contrasts, and blocking structures that are not standard in existing single-cell pseudobulk workflows and are rarely implemented in published analyses. Moreover, these tools produce gene-level regression coefficients from a pooled model rather than participant-level effect estimates, making it difficult to inspect, visualize, or perform sensitivity analyses on the within-participant contrasts that are the natural estimand in repeated-measures intervention studies. Practical tradeoffs among scalability, stability, and interpretability further constrain their use in the small-sample, multi-time-point settings typical of clinical single-cell studies^28^.

Difference-in-differences (DiD) estimation offers a natural framework for this inferential structure. Widely used in econometrics and policy evaluation for over three decades^29–31^, DiD estimates between-group differences in within-unit change over time while controlling for baseline differences and temporal effects shared across groups. When applied at the level of biological replication, the estimand is the difference between groups in their within-participant pre-to-post change, an interpretable participant-level contrast that is well suited to longitudinal intervention studies^32,33^. Yet despite its rapid adoption in epidemiology and health policy evaluations^34^, DiD has not been adapted as a participant-level inferential framework for longitudinal single-cell transcriptomics. Existing pseudobulk and mixed-effects workflows can in principle accommodate longitudinal interaction terms through manual design specification. However, they are not organized around design-specific participant-level estimands, do not naturally expose within-participant contrasts for inspection and sensitivity analysis, and do not provide a unified workflow that links study architecture to inference, diagnostics, and interpretation. This gap leaves an important class of questions, whether treatments or clinically defined groups exhibit distinct transcriptional trajectories over time, without a dedicated inferential framework for single-cell data.

Here, we present *sctrial*, a design-aware Python package for longitudinal single-cell inference that organizes analysis around the experimental design and the participant-level estimand it implies. Specifically, *sctrial* formalizes DiD as the default participant-level estimand for two-group longitudinal studies and extends the same design-aware logic to paired pre/post studies and cross-sectional comparisons. The package couples these estimands to inference procedures tailored to small effective sample sizes, integrating pseudobulk summaries with cluster-aware uncertainty estimates, resampling-based sensitivity analyses, and pathway-aware effect ranking.

We applied *sctrial* to five independent datasets spanning melanoma immune checkpoint blockade (ICB)^35^, COVID-19 severity^36^, BNT162b2 vaccination^37^, acute myeloid leukemia (AML) chemotherapy^38^, and CAR-T therapy^39^ to evaluate how these design-specific analyses behave across distinct immunological and clinical contexts. Across these applications, *sctrial* attenuated significance inflation driven by pseudoreplication, produced interpretable effect estimates, and revealed context-dependent immune programs whose magnitude and direction varied across intervention settings, disease states, and follow-up windows. Together, these analyses establish *sctrial* as a framework for repeated-measures single-cell studies, particularly in clinical-trial and translational settings where the key objective is to determine whether treatments, disease states, or response groups induce distinct within-participant transcriptional trajectories over time.

## RESULTS

### Cell-Level Analysis Inflates Apparent Significance Relative to Participant-Level Inference

We first establish the inferential framework implemented in *sctrial* and the design-specific estimands it returns (**Figure 1**). Given an AnnData object and study-design specification, *sctrial* performs participant-by-visit pseudobulk aggregation, verifies analyzable repeated measures, and returns design-aligned participant-level effects: DiD for two-group longitudinal studies, paired change for single-arm pre/post studies, and standardized between-group contrasts for cross-sectional comparisons.

**Figure 1.**
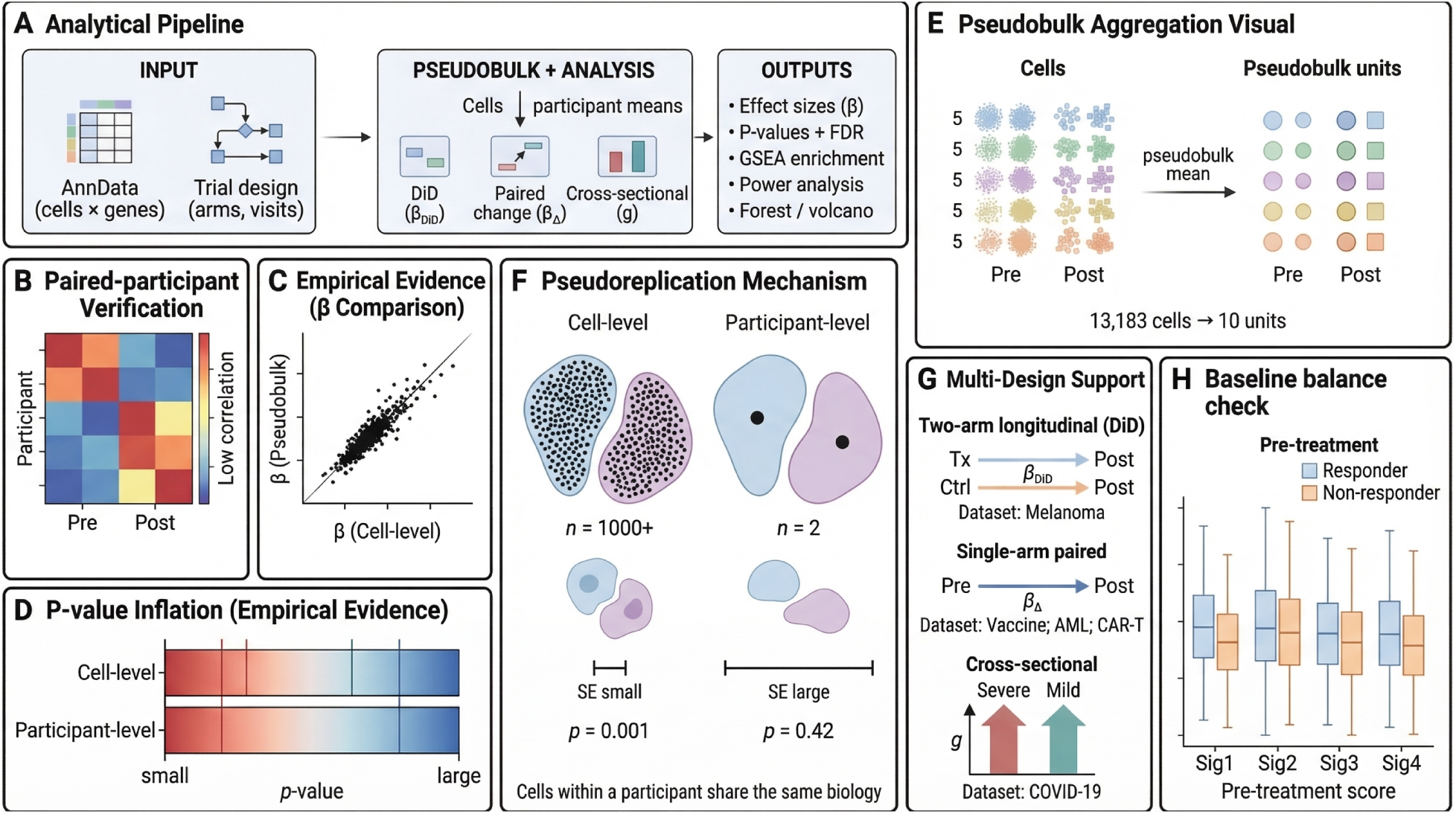
Overview of the sctrial framework for participant-level inference in longitudinal single-cell studies. **(A)** sctrial accepts an AnnData object and a trial design specification (treatment arms, visits) as input, performs pseudobulk aggregation to participant-level means, and supports three analysis modes: difference-in-differences (β_DiD_) for two-arm longitudinal designs, paired within-participant change (β_Δ_) for single-arm studies, and cross-sectional effect sizes (Hedges’ g) for between-group comparisons. Outputs include effect sizes with confidence intervals (CIs), false discovery rate (FDR)-corrected p-values, gene set enrichment analysis (GSEA), power calculations, and visualizations. **(B)** Heatmap illustrating within-participant correlation across time points. **(C)** Scatter plot comparing cell-level and pseudobulk (participant-level) β estimates. **(D)** Conceptual comparison of –log₁₀(p) between cell-level and participant-level analyses, showing that cell-level inference produces systematically smaller p-values due to pseudoreplication. **(E)** Pseudobulk aggregation: cells from each participant and time point (colored by participant identity) are collapsed into a single pseudobulk mean, reducing 13,183 cells to 10 participant-level units (using the melanoma dataset as example). **(F)** The pseudoreplication mechanism: at the cell level (left), thousands of non-independent observations yield artificially small standard errors and inflated significance (p = 0.001). At the participant level (right), the effective sample size reflects the true number of biological replicates (n = 2 per group), producing appropriately large standard errors and calibrated p-values (p = 0.42). **(G)** sctrial accommodates three study architectures: two-arm longitudinal DiD (melanoma; responder vs. non-responder, pre vs. post treatment), single-arm paired designs (vaccine, AML, CAR-T; within-participant pre–post change), and cross sectional between-group comparisons (COVID-19; severe vs. mild). **(H)** Box plots comparing pre-treatment signature scores between responders and non-responders across representative signatures, assessing whether groups are comparable prior to intervention.

To evaluate the practical consequences of the inferential unit choice, we assembled five publicly available scRNA-seq datasets spanning melanoma immunotherapy^35^, COVID-19 severity^36^, BNT162b2 vaccination^37^, AML chemotherapy^38^, and CAR-T therapy^39^ (**Table 1**). Dataset preprocessing, quality control, and annotation procedures are described in **Methods** and **Supplementary Figures 1–2**. We then used the melanoma immunotherapy cohort as a benchmark to compare cell-level and participant-level inference under the same biological contrast.

**Table 1.** Summary of datasets and analyzable subsets used in sctrial analyses. ICB: immune checkpoint blockade; QC: quality control; DFO: days from onset; AML: acute myeloid leukemia.

| Dataset | Data Source | Study Design | Total Participants (N paired) | N Per-arm | Timepoints | Analysis Type in scRNA |
| --- | --- | --- | --- | --- | --- | --- |
| **Melanoma ICB** | GSE120575 | Two-arm parallel (Responder vs Non-responder) Pre/Post ICB | 25 (10 paired; both Pre + Post) | 3 Responders<br>7 Non-responders (after QC filtering) | Pre-ICB<br>Post-ICB | Difference-in-Differences (DiD) |
| **COVID-19 Severity** | E-MTAB-10026 | Cross-sectional (Severe vs Mild) Multiple DFO bins | 143 (16; in DFO 8–14 analysis window) | 8 Severe<br>8 Mild (DFO 8–14 bin) | DFO 0–7<br>DFO 8–14<br>DFO 15+ | Between-arm Hedges' g |
| **BNT162B2 Vaccination** | GSE171964 | Single-arm paired Pre/Post vaccination | 21 (6; paired Day 0 + Day 7) | All treated (single-arm) | Day 0<br>Day 7 | Within-arm paired $\Delta$ |
| **AML Chemotherapy** | GSE116256 | Longitudinal Diagnosis - Post-chemo (Treatment arm only) | 21 (11 paired; Treatment arm with Pre + Post) | All treated (single-arm; healthy controls excluded) | Diagnosis<br>Day 15–18<br>Day 31–41 | Within-arm paired $\Delta$ |
| **CAR-T Therapy** | GSE290722 | Single-arm paired Pre/Post CAR-T infusion | 50 (31 paired; with both Pre + Post) | All treated (single-arm) | Day -28<br>Day +28 (+180, +365) | Within-arm paired $\Delta$ |

The melanoma dataset included 25 participants, of whom 10 had paired pre-and post-treatment samples suitable for longitudinal analysis (3 responders and 7 non-responders; **Figure 2A**). We curated 12 immune signatures representing major axes of anti-tumor immunity, including cytotoxicity, exhaustion, interferon signaling, inflammation, and antigen presentation (**Supplementary Table 1**). For each signature, we estimated the same responder-versus-non-responder difference in pre-to-post change, while varying only the unit treated as statistically independent: individual cells for the cell-level analysis (n = 13,183 cells) versus participant-level pseudobulk summaries for the DiD analysis (n = 10 paired participants).

**Figure 2.**
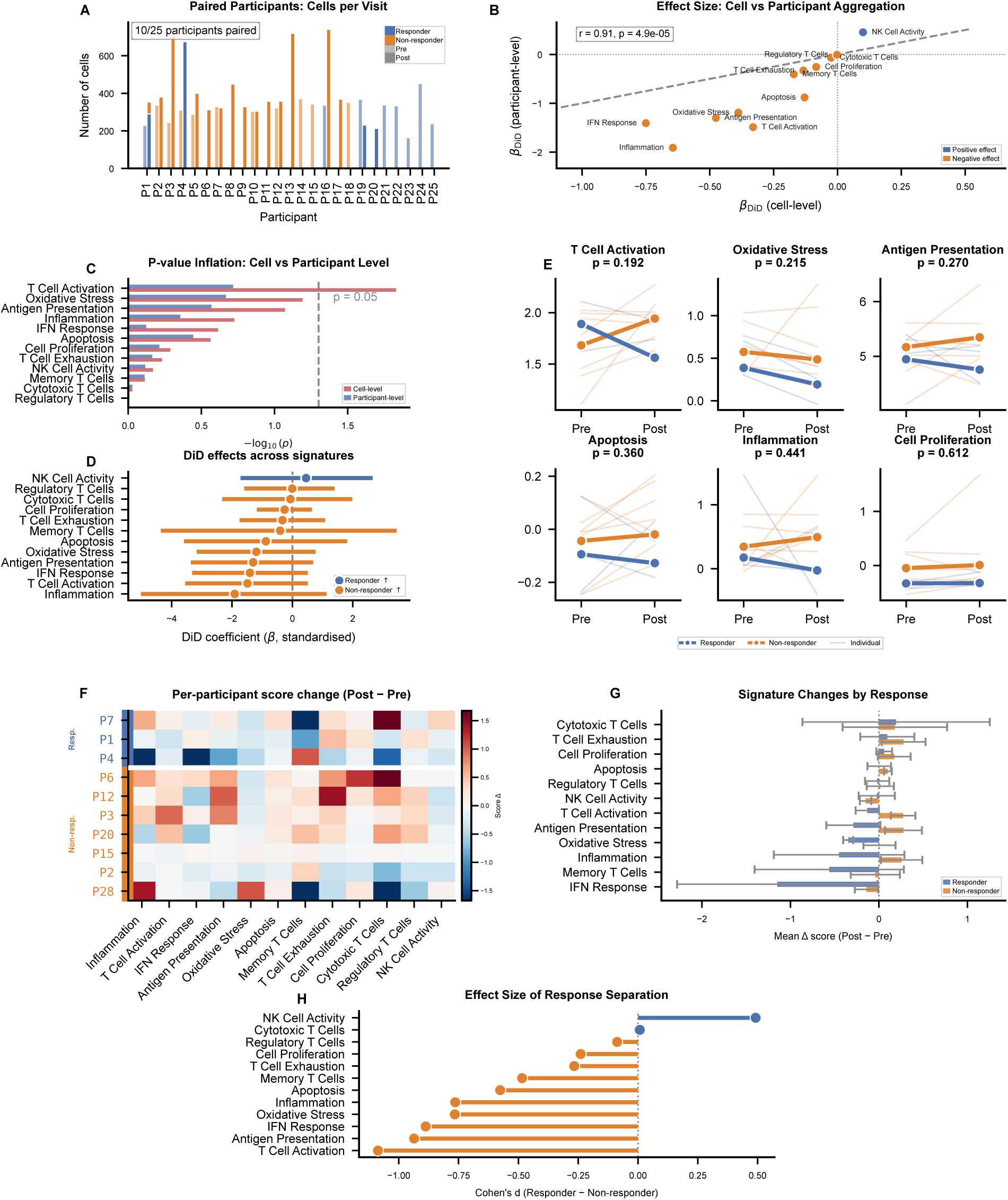
Participant-level analysis of the melanoma immunotherapy cohort. **(A)** Grouped bar chart showing the number of cells per participant stratified by visit and response status. **(B)** Scatter plot of cell-level versus the participant-level DiD coefficients for each immune signature. Pearson correlation coefficient and p-value are shown. **(C)** Bar charts comparing –log₁₀(p) from cell-level and participant-level analyses. **(D)** Forest plot of signatures’ DiD coefficient with 95% bootstrap-t confidence intervals. **(E)** Line plots for the six most significant signatures showing individual participant trajectories and group means from pre-to post-treatment, stratified by response. **(F)** Heatmap of per-participant change (Δ = Post − Pre) for each immune signature. **(G)** Horizontal bars showing the mean post-minus-pre change (Δ) for responders and non-responders with standard error bars, for each signature. **(H)** Lollipop chart of Cohen’s d (responder − non-responder) for each signature, sorted by magnitude.

The two approaches yielded broadly concordant effect directions across signatures (Pearson r = 0.914, p = 4.9 × 10⁻⁵; **Figure 2B**), but agreement in direction did not translate into agreement in statistical inference. When we compared nominal p-values (**Figure 2C**), the cell-level analysis identified T cell activation as significant (p = 0.014), with oxidative stress (p = 0.060) and antigen presentation (p = 0.079) nominally non-significant at α = 0.05. The participant-level analysis yielded p = 0.113 for T cell activation, and no signature remained significant after accounting for the effective sample size of 10 paired participants (all FDR > 0.47; **Supplementary Table 2**). This divergence reflects the fact that cell-level analysis treats 13,183 correlated cells as independent observations, producing standard errors that are artificially small relative to an analysis that respects participant-level replication. Several signatures also showed larger absolute effect estimates at the participant level than at the cell level, including inflammatory response (*β_DiD_* = −1.91), T cell activation (*β_DiD_* = −1.49), and IFN response (*β_DiD_* = −1.41), all indicating greater pre-to-post increases in non-responders than in responders (**Figure 2B**). Thus, even when cell-level and participant-level analyses agree qualitatively in direction, they can yield materially different uncertainty estimates, significance assessments, and effect magnitudes.

### sctrial Characterizes Response-Associated Participant-Level Immune Signature Trajectories in Melanoma

Having established that cell-level testing overstates statistical evidence in the melanoma cohort, we next used *sctrial* to characterize participant-level immune signature trajectories associated with ICB. For each of the 12 curated immune signatures, we estimated a DiD effect using participant-by-timepoint pseudobulk summaries from the 10 paired participants, with uncertainty quantified by 1,000 wild cluster bootstrap resamples and p-values adjusted using the Benjamini-Hochberg procedure.

Across the 12 signatures, the largest DiD estimates were negative, indicating greater pre-to-post increases in non-responders than in responders (**Figure 2D**). The top-ranked signatures all showed wide bootstrap-derived confidence intervals, including inflammation (*β_DiD_* = −1.91, 95% CI: −4.30 - 0.48), T cell activation (*β_DiD_* = −1.49, 95% CI: −3.13 - 0.15), IFN response (*β_DiD_* = −1.41, 95% CI: −3.02 - 0.21), and antigen presentation (*β_DiD_* = −1.30, 95% CI: −2.93 - 0.33). NK-cell activity was the only signature with a positive DiD estimate (*β_DiD_* = 0.45, 95% CI: −1.44 - 2.35).

Participant-level trajectories for the six signatures with the smallest nominal p-values illustrated the basis of this imprecision (**Figure 2E**). Although mean trends separated between response groups for T cell activation (bootstrap p = 0.192), oxidative stress (p = 0.215), and antigen presentation (p = 0.270), individual trajectories showed substantial between-participant heterogeneity within each response category (**Figure 2F**). Group-level Δ scores (post minus pre) supported this directional separation: T cell activation decreased on average in responders (mean Δ = −0.14) but increased in non-responders (mean Δ = 0.43), and antigen presentation showed a similar pattern (responder Δ = −0.22, non-responder Δ = 0.37; **Figure 2G**). These results indicate that dominant limitation in this cohort is imprecision arising from marked inter-participant variability, rather than lack of directional separation in the group means.

To facilitate cross-cohort comparison on a standardized scale, we also computed Hedges’ g from participant-level Δ scores (**Figure 2H**). T cell activation showed the largest negative standardized effect (d = −1.05), followed by antigen presentation (d = −1.00), IFN response (d = −0.92), oxidative stress (d = −0.78), and inflammation (d = −0.72), whereas NK-cell activity had the largest positive effect size (d = 0.50). These standardized effects were directionally consistent with the DiD estimates and provide a scale for comparing signature shifts across cohorts.

### Sensitivity Analyses Characterize the Stability and Uncertainty of Participant-Level Inference

We next evaluated the robustness of participant-level inference with sensitivity analyses targeting uncertainty estimation, preprocessing dependence, and the influence of individual participants. Because standard cluster-robust inference can be anti-conservative when the number of clusters is small^40^, we compared analytical cluster-robust standard errors (SEs) for the DiD coefficients with SEs estimated from 1,000 wild cluster bootstrap resamples. Residual plots and additional model diagnostics for all datasets are shown in **Supplementary Figure 3A–I**. Across the 12 immune signatures, analytical and bootstrap SEs were broadly concordant (r = 0.76, p = 4.3 × 10⁻³; **Figure 3A**; **Supplementary Figure 4A**), with several signatures lying close to the identity line. However, inflammation (analytical SE = 1.22, bootstrap SE = 1.90) and IFN response (analytical SE = 0.82, bootstrap SE = 1.51) showed substantially larger bootstrap SEs, indicating that analytical inference can underestimate uncertainty for some signatures in small-sample settings.

**Figure 3.**
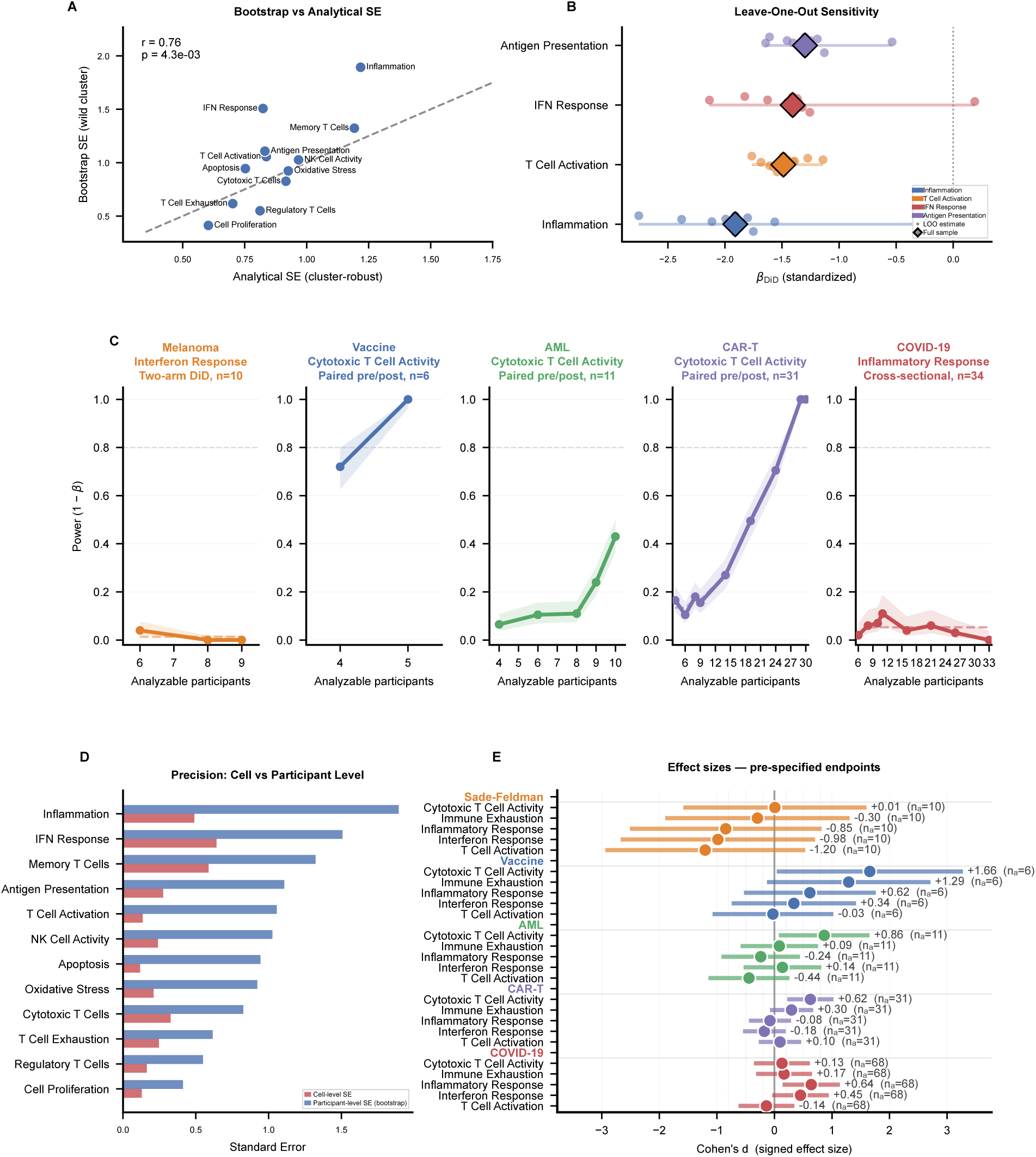
Robustness, benchmarking, and cross-dataset effect sizes. **(A)** Scatter plot comparing analytical (cluster-robust) standard errors (x-axis) with wild cluster bootstrap standard errors (y-axis) across immune signatures in the melanoma cohort. Pearson correlation coefficient is shown; the dashed line indicates perfect agreement. **(B)** Range plot showing the distribution of DiD coefficients when each participant is systematically excluded, for the four most influential signatures. Diamonds indicate the full sample estimate; circles indicate leave-one-out estimates. **(C)** Empirical power curves; for each dataset, the relationship between the number of analyzable participants (x-axis) and statistical power (y-axis) was estimated by subsampling participants and computing the fraction of iterations in which the pre-specified endpoint reached significance. Shaded ribbons indicate Wilson confidence intervals; dashed lines show isotonic regression fits. The horizontal dashed line marks 80% power. **(D)** Bar plot comparing cell-level standard errors with participant-level bootstrap standard errors. **(E)** Forest plot showing Cohen’s d with 95% confidence intervals for the primary endpoint in each dataset, grouped by study. Sample sizes per dataset are annotated. Vertical dashed lines indicate conventional effect-size thresholds (±0.2, ±0.5, ±0.8).

Effect estimates were also stable across alternative preprocessing choices. Concordance between standardized and unstandardized effects is shown in **Supplementary Figure 4B**. Rank-order concordance of DiD coefficients remained high under alternative normalization and aggregation strategies (**Supplementary Figure 4C–D**), and cell-type-stratified DiD coefficients showed consistent rank ordering across preprocessing variants (**Supplementary Figure 4E–F**). Together, these analyses indicate that the main participant-level trends were not driven by a single normalization or aggregation choice.

We then assessed the influence of individual participants by repeating the DiD analysis in a leave-one-out framework for the four signatures with the largest absolute DiD effects (**Figure 3B**). T cell activation and antigen presentation were relatively stable across exclusions, indicating that their estimated effects were not driven by any single participant. In contrast, inflammation and IFN response were more sensitive to individual omissions, with some leave-one-out estimates shifting materially toward or away from zero (**Supplementary Figure 3C**). Maximum leave-one-out deviations across features and datasets are summarized in **Supplementary Figure 4G**. These results show that participant-level effect estimates can distinguish between robust directional trends and signatures whose apparent magnitude is more dependent on individual participants.

### Simulation Benchmark Reveals Calibration Tradeoffs Across Methods

To evaluate statistical behavior under controlled conditions, we conducted a Monte Carlo simulation using a hierarchical gamma-Poisson generative model calibrated from the melanoma immunotherapy dataset (see **Methods**). We compared *sctrial*’s first-difference DiD estimator with dreamlet^41^, NEBULA^27^, and a nonparametric comparator applied to participant-level change scores, across sample sizes of n = 8, 12, 20, 40, and 60 participants per arm and true effect sizes of β = 0, 0.2, 0.5, and 1.0. Under pure-null conditions (β = 0), all four methods maintained empirical Type I error rates near the nominal 5% level across sample sizes (**Supplementary Figure 4I**), with genomic inflation factors close to unity (**Supplementary Figure 4J**) and null-gene p-value distributions tracking the expected uniform diagonal (**Supplementary Figure 4K**). Under non-null conditions, all methods showed similar power on truly affected genes, approaching saturation at moderate effect sizes for n ≥ 20 (**Supplementary Figure 4H**). Similar power patterns were observed under the single-arm paired design (**Supplementary Figure 4M**).

The principal differences emerged in mixed-signal panels containing both affected and unaffected genes. Under the evaluated pipeline configurations, dreamlet showed substantial false positive inflation among null genes, rising from approximately 20% at β = 0.2 to over 80% at β = 1.0 (**Supplementary Figure 4L**), whereas NEBULA showed more modest inflation under the same conditions. By contrast, *sctrial* and the nonparametric change-score comparator remained close to nominal across the evaluated settings. Consistent with this pattern, null-gene p-value calibration was stable for all four methods under pure-null scenarios but deteriorated markedly for dreamlet under mixed-signal conditions (**Supplementary Figure 4K**). Computationally, *sctrial* and the nonparametric comparator completed each iteration in approximately 1 millisecond, compared with 1–2 seconds for dreamlet and NEBULA (**Supplementary Figure 4N**).

To complement the simulation benchmark with data-conditional estimates from real studies, we next computed resampling-based power curves across all five datasets using a pre-declared primary endpoint for each (**Figure 3C**). Power was defined as the proportion of 200 participant-level subsampling iterations yielding p < 0.05, with isotonic regression enforcing monotonicity and Wilson confidence intervals summarizing uncertainty. In this observed subsampling analysis, the BNT162b2 vaccination dataset (cytotoxic T-cell activity, n = 6) showed near-maximal empirical power, consistent with its large observed effect size (d = 1.66), whereas the AML dataset (n = 11) reached approximately 45% power at its maximum analyzable sample size. The remaining cohorts, including melanoma (n = 10), CAR-T (n = 31), and COVID-19 (n = 34), remained below 25% power across evaluated subsample sizes. Although these estimates are conditional on the observed datasets and endpoints, they suggest that many currently available longitudinal single-cell studies are underpowered for participant-level inference at conventional significance thresholds.

Finally, comparison of cell-level and participant-level standard errors across all five cohorts confirmed that cell-level SEs were uniformly smaller (**Figure 3D**; **Supplementary Figure 3J**), with the largest discrepancies observed for T cell activation (cell-level SE = 0.14 vs. participant-level bootstrap SE = 1.05) and inflammation (0.50 vs. 1.90) in the melanoma cohort. These differences directly quantify how cell-level inference overstates precision when within-participant correlation is ignored^42^.

To place these findings in a broader context, we compared standardized participant-level immune effects across all five cohorts by computing signed Cohen’s d values for five pre-specified immune signatures (**Figure 3E**). Cytotoxic T cell activity showed the largest positive effects in the BNT162b2 vaccine (d = 1.66, n = 6), AML (d = 0.90, n = 11), and CAR-T (d = 0.49, n = 31) datasets, whereas inflammatory response was the dominant positive effect in COVID-19 (d = 0.64, n = 34). In the melanoma cohort, most signatures were negative, with T cell activation showing the strongest negative effect (d = −1.20, n = 10), indicating greater pre-to-post increases in non-responders than responders.

### Pathway Analysis Reveals Cohort-Specific and Shared Transcriptional Trends

To identify broader transcriptional programs beyond curated signatures, we performed pre-ranked gene set enrichment analysis (GSEA)^43^ on participant-level gene effects within each cohort using MSigDB Hallmark^44^, KEGG^45^, Reactome^46^, GO Biological Process^47^, and WikiPathways^48^ collections (**Figure 4**; **Supplementary Table 3**). Genes were ranked within each cohort by the design-specific participant-level effect estimate, using *β_DiD_* for two-group longitudinal analyses and paired Δ for single-arm analyses, and enrichment was summarized by the normalized enrichment score (NES).

**Figure 4.**
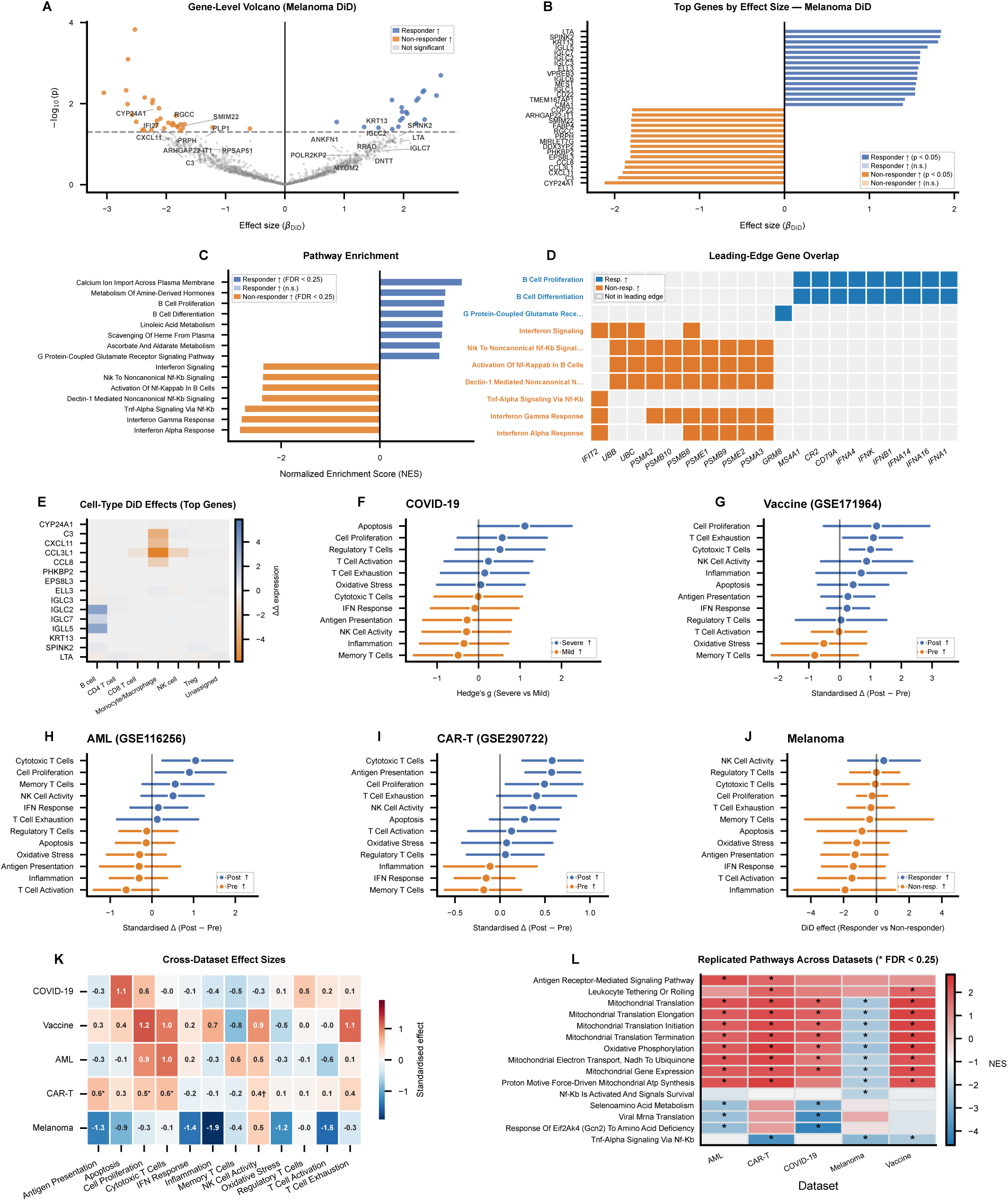
Gene-level and pathway-level biological discovery. **(A)** Gene-level volcano plot in the melanoma cohort. The x-axis shows the DiD coefficient and the y-axis shows –log₁₀(p). Genes are colored by direction and significance: blue (responder-associated, p < 0.05), orange (non-responder-associated, p < 0.05), and grey (not significant). Top protein-coding genes in each direction are labelled. **(B)** Waterfall bar chart of the genes with the largest positive and negative β_DiD_ values (15 genes each) in the melanoma cohort. **(C)** Bar chart showing normalized enrichment scores (NES) for the most strongly enriched pathways across MSigDB Hallmark, Reactome, KEGG, GO Biological Process, and WikiPathways libraries. **(D)** Heatmap of shared leading-edge genes across the 10 most strongly enriched pathways, and colored by response status. Columns (genes) are hierarchically clustered by Jaccard distance. **(E)** Heatmap of mean DiD-like effects for the top 15 genes (by β_DiD_) across annotated cell types. **(F)** COVID-19 severity (n = 16 at DFO 8–14 days). Forest plot of Hedges’ g (severe minus mild) for each immune signature computed as a cross-sectional between-group comparison. **(G)** BNT162b2 vaccination (GSE171964, n = 6). Within-participant standardized change (Δ, day 7 minus day 0) with bootstrap confidence intervals. **(H)** AML chemotherapy (GSE116256, n = 11 paired). Within-participant standardized change (post-treatment minus diagnosis) for the treatment arm. **(I)** CAR-T therapy (GSE290722, n = 31 paired). Within-participant standardized change (day +28 minus day −28). **(J)** Melanoma immunotherapy (Sade-Feldman, n = 10 paired). DiD coefficients (responder versus non-responder) with bootstrap confidence intervals. **(K)** Heatmap of standardized effect sizes of immune signatures (rows) across datasets (columns). Note that the estimand differs by design (β_DiD_ for melanoma, Δ for single-arm cohorts, Hedges’ g for COVID-19). **(L)** Heatmap of NES of overlapping pathways across datasets. Shown are pathways present in at least two datasets. Asterisks denote significant pathways (FDR < 0.25).

In the melanoma cohort, participant-level DiD analysis showed a modest overall skew toward non-responder-associated transcriptional effects (**Figure 4A**). Among statistically significant genes, more had negative than positive *β_DiD_* values, but only two genes remained significant after transcriptome-wide FDR correction: RP11-232M24.1 (*β_DiD_* = 2.67, FDR = 0.004) and FDCSP (*β_DiD_* = 2.66, FDR = 0.005), both in the responder direction (**Figure 4B**; **Supplementary Table 6**). Among the strongest responder-direction genes, immunoglobulin-related transcripts were prominent, including IGLC2, IGLC3, IGLC7, IGLL5, IGLC6, and IGLC1, together with LTA, SPINK2, and KRT13. By contrast, the strongest non-responder-direction genes included CYP24A1, C3, CXCL11, CCL3L1, and CCL8. Taken together, the gene-level landscape suggests a responder-associated humoral/B-lineage program and a non-responder-associated inflammatory chemokine/complement program.

GSEA of the ranked melanoma gene list showed the strongest enrichment in the non-responder direction (**Figure 4C**; **Supplementary Table 3**). Among the most strongly enriched Hallmark pathways were interferon alpha response (NES = −2.84, FDR q < 0.001), interferon gamma response (NES = −2.81, q < 0.001), TNFα signaling via NF-κB (NES = −2.74, q < 0.001), and oxidative phosphorylation (NES = −2.51, q < 0.001). Related Reactome terms, including interferon signaling and multiple NF-κB-associated modules, were enriched in the same direction. Together, these results support a coordinated non-responder-associated inflammatory and interferon-linked transcriptional program. Several responder-direction pathways, including calcium ion import across the plasma membrane, B cell proliferation, B cell differentiation, and linoleic acid metabolism, also showed positive NES values. However, these pathways did not survive multiple testing correction across the tested pathway universe, consistent with the limited power afforded by 10 paired participants to detect enrichment in the responder direction.

Leading-edge analysis of the top enriched pathways identified the genes contributing most strongly to these signals (**Figure 4D**). On the responder side, B cell proliferation and B cell differentiation shared leading edge genes that include CD79A and MS4A1, together with several type I interferon family members, consistent with prior evidence linking type I interferon signaling to B cell activation and differentiation^49^. On the non-responder side, multiple NF-κB-and interferon-associated pathways shared proteasome-related genes, including PSMB10, PSMB8, PSMA3, PSME1, PSMB9, and PSME2, and the interferon-stimulated gene IFIT2 recurred across several leading edges. This overlap supports a coordinated interferon-linked inflammatory program involving immunoproteasome components, consistent with prior evidence that interferons induce immunoproteasome subunits participating in inflammatory signaling^50^.

Cell type-resolved DiD analysis further contextualized these effects across specific populations (**Figure 4E**). Immunoglobulin-related responder-direction genes, including IGLC2, IGLC3, IGLC7, and IGLL5, showed their strongest effects in B cells and plasma cells. In contrast, non-responder-direction genes such as CCL3L1, CXCL11, CCL8, and C3 showed their strongest effects in monocyte/macrophage populations. These cell type-resolved patterns reinforce the distinction between a humoral-associated responder program and a myeloid-associated inflammatory program in non-responders.

### Application Across Distinct Longitudinal and Quasi-Longitudinal Cohorts

We next applied *sctrial* to four additional datasets spanning COVID-19 severity^36^, BNT162b2 vaccination^37^, AML chemotherapy^38^, and CAR-T therapy^39^. Because the underlying study designs differed across cohorts, each dataset was summarized using the participant-level effect metric appropriate to its design: Hedges’ g for the cross-sectional COVID-19 comparison, standardized within-participant change (post minus pre) for the single-arm BNT162b2, AML, and CAR-T cohorts, and *β_DiD_* for the two-group longitudinal melanoma analysis. These quantities were therefore used for within-cohort inference and descriptive cross-cohort comparison, rather than as identical estimands on a common causal scale. Gene set score distributions for all cohorts are shown in **Supplementary Figure 5A**.

Within each cohort, participant-level analysis identified distinct immune signature profiles (**Figures 4F–J; Supplementary Figure 5A**). In the COVID-19 severity cohort^36^, apoptosis showed the largest severity-associated effect (Hedges’ g = 1.12, p = 0.043), followed by cell proliferation (g = 0.57) and regulatory T cells (g = 0.52), whereas memory T cells showed a negative effect (g = −0.49). No signature survived multiple testing correction (**Figure 4F**).

In the BNT162b2 vaccination cohort (n = 6 paired participants)^37^, the largest post-vaccination increases were observed for cell proliferation (Δ = 1.2), T cell exhaustion (Δ = 1.1), cytotoxic T cells (Δ = 1.0), NK-cell activity (Δ = 0.9), and inflammation (Δ = 0.7), together with decrease in memory T cells (Δ = −0.8) and oxidative stress (Δ = −0.5) (**Figure 4G**). In AML (n= 11)^38^, cytotoxic T cells showed the largest positive post-treatment shift (Δ = 1.1), while apoptosis (Δ = −0.7) and T cell activation (Δ = −0.7) decreased (**Figure 4H**). The CAR-T cohort (n= 31)^39^ showed more modest post-infusion changes, with cell proliferation (Δ = 0.6), antigen presentation (Δ = 0.5), and cytotoxic T cells (Δ = 0.4) showing the largest positive effects (**Figure 4I**).

A cross-dataset heatmap of standardized effect summaries revealed both recurring and context-specific immune patterns (**Figure 4K**; **Supplementary Figure 5B**). Cytotoxic T cell-activity was upregulated in the BNT162b2, AML, and CAR-T cohorts, whereas other signatures changed direction across settings. For example, T cell activation was negative in melanoma and AML but near zero in the BNT162b2 and CAR-T datasets, and inflammation was negative in melanoma and AML but positive in the BNT162b2 cohort (**Supplementary Figure 5C–G**). These comparisons indicate that some immune programs show broadly concordant directional behavior across intervention settings, whereas others are strongly context-dependent.

Cohort-level GSEA identified pathway signals that recurred across datasets at FDR < 0.25 (**Figure 4L**; **Supplementary Figure 5H**; **Supplementary Table 3**). E2F targets were positively enriched in both AML and melanoma, while mitochondrial translation-related pathways were positively enriched in CAR-T and COVID-19. TNFα signaling via NF-κB showed its strongest negative enrichment in melanoma, with related negative enrichment also observed in AML. Together, these pathway-level comparisons suggest that cell-cycle and mitochondrial programs recur across multiple immune perturbation settings, whereas inflammatory programs are more strongly context dependent.

### Permutation Analysis and Temporal Stratification Clarify Participant-Level Effects

To assess whether the observed melanoma effects exceeded chance expectation, we performed a participant-level permutation analysis in which response labels were permuted across participants while preserving each participant’s paired pre/post structure, and the DiD statistic was recomputed across 1,000 permutations. IFN response yielded the smallest empirical permutation p-value (p = 0.092), followed by T cell activation (p = 0.101) and antigen presentation (p = 0.146) (**Figure 5A**; **Supplementary Table 4**). For IFN response, the observed DiD estimate (β = −1.41) lay toward the lower tail of the permutation distribution, but no signature reached statistical significance (**Figure 5B**). These results were directionally consistent with the parametric and bootstrap-based analyses while underscoring the limited resolution of permutation-based inference in a cohort of 10 paired participants (**Supplementary Table 5**).

**Figure 5.**
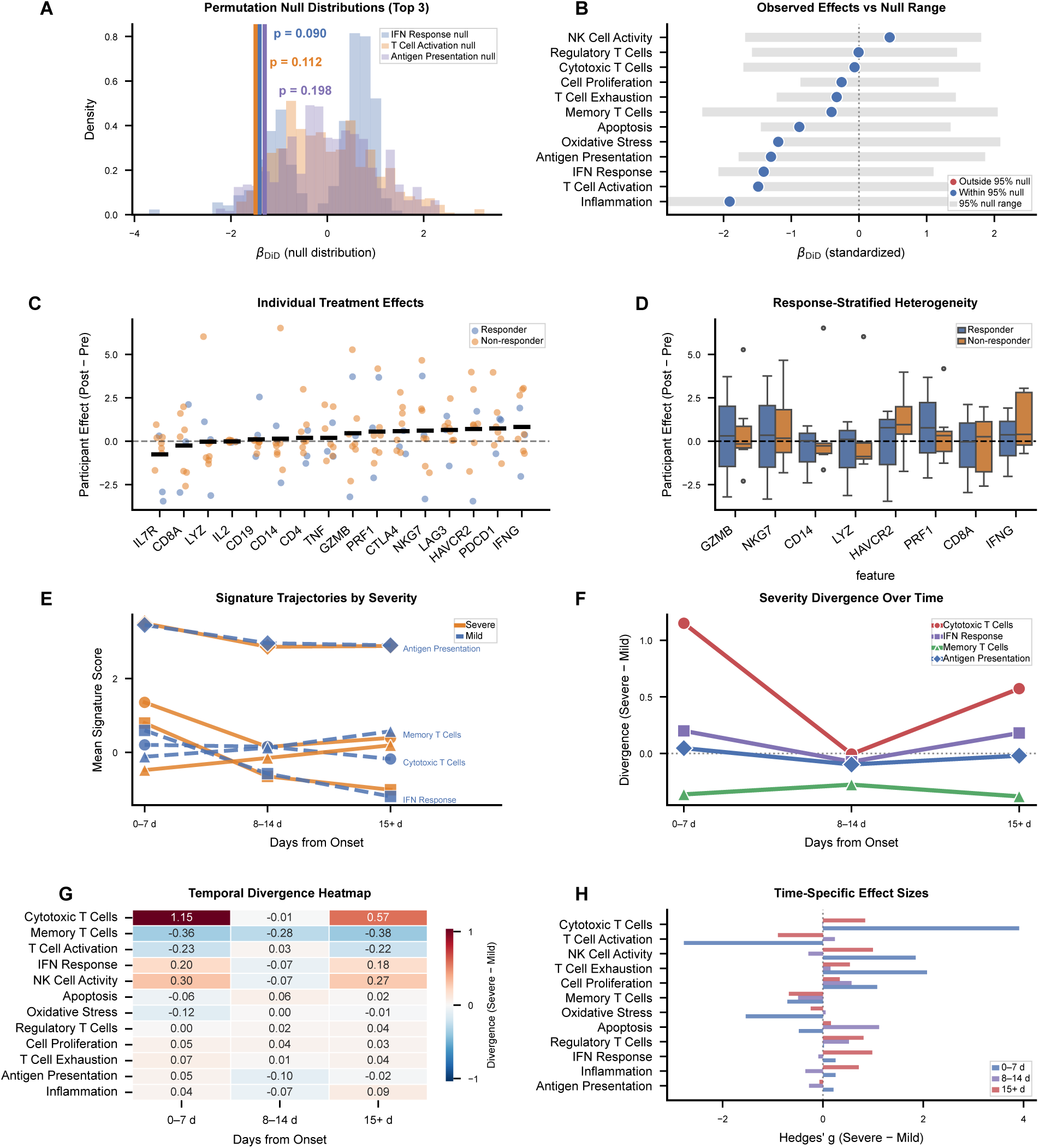
Permutation-based validation, participant-level heterogeneity, and temporal severity dynamics. **(A)** Histograms of DiD coefficients from 1,000 participant-level label permutations for the three signatures with the smallest permutation p-values in the melanoma cohort. Vertical lines indicate the observed effects. **(B)** Observed effects versus permutation null range. For each signature, the horizontal grey bar spans the 2.5th–97.5th percentiles of the permutation distribution. Points falling outside the grey band (red) indicate effects exceeding the 95% permutation null; points within (blue) indicate effects consistent with label shuffling. **(C)** Strip plot of per-participant post-minus-pre changes across 16 immune-related genes, stratified by clinical response. Horizontal lines indicate group means. **(D)** Box plots of participant-level effects for the eight genes with the highest inter-participant standard deviation, comparing responders and non-responders. **(E–F)** Temporal severity dynamics in the COVID-19 severity cohort. **(E)** Mean signature scores for severe and mild groups across three temporal bins (DFO 0–7, 8–14, and 15+ days) for four representative signatures. **(F)** Severity divergence (severe minus mild) over time for the same four signatures. **(G)** Temporal divergence heatmap. All 12 immune signatures (rows) × three DFO bins (columns), with color encoding the severe-minus-mild difference. **(H)** Grouped horizontal bar chart of Hedges’ g (severe minus mild) for each signature within each temporal bin.

We next examined participant-level post-treatment changes in 16 immune-related genes spanning T cell markers (CD8A, CD4, IL7R), exhaustion-associated receptors (PDCD1, HAVCR2, LAG3, CTLA4), cytotoxic effectors (GZMB, PRF1, NKG7), cytokines (IFNG, TNF, IL2), and myeloid/B-cell markers (CD14, LYZ, CD19) stratified by response status (**Figure 5C**; **Supplementary Figure 6A-C**). For several genes, between-participant variation was comparable to or larger than the mean group-level shift, with substantial overlap between responders and non-responders (**Figure 5D; Supplementary Figure 6D–E**). IL7R tended to decrease in responders while remaining closer to baseline in non-responders, LYZ shifted in opposite directions across the two groups, and CTLA4 increases were more prominent among non-responders (**Supplementary Figure 6F–J**). These patterns illustrate how participant-level analyses makes heterogeneity in treatment response directly visible, rather than absorbing it into a pooled residual variance term.

To examine whether participant-level analysis can also resolve time-varying effects beyond the melanoma benchmark, we next analyzed the COVID-19 cohort across the disease course. Samples were stratified into three bins based on days from symptom onset (DFO: 0–7, 8–14, and 15+ days), and severity divergence was quantified as the difference between severe and mild participant-level means for each signature (**Figure 5E–F**). Cytotoxic T cell activity displayed the largest early divergence, minimal intermediate separation, and renewed late divergence; IFN response showed a similar non-monotonic pattern. Memory T cell scores remained consistently lower in severe disease across all three phases. Across all immune programs (**Figure 5G–H**), cytotoxic T cell and NK cell-associated scores showed the strongest early severity separation, whereas apoptosis, oxidative stress, regulatory T cells, and cell proliferation remained comparatively close to zero throughout. These results are consistent with the findings of Stephenson et al. that immune-state differences in COVID-19 evolve over the disease course^36^, and they illustrate how temporally stratified participant-level analysis can resolve dynamics that would be obscured by a single pooled cross sectional contrast.

## DISCUSSION

The growing use of scRNA-seq in longitudinal, repeated-sampling, and interventional studies has created a need for analytical strategies that go beyond cross-sectional differential expression and instead align inference with both biological replication and study design^2^. In multi-participant single-cell datasets, two challenges are central: avoiding pseudoreplication by treating the participant rather than the cell as the inferential unit, and matching the estimand to the question posed by the study design. Two-group treatment-response studies, paired pre/post interventions, and cross-sectional severity comparisons target different participant-level quantities, and collapsing them into a single testing framework obscures both interpretation and uncertainty. *sctrial* addresses this by linking each study architecture to a design-specific participant-level estimand and coupling these estimands to small-cluster-aware uncertainty quantification, participant-level diagnostics, and pathway-level interpretation.

Our results illustrate the practical consequences of this formulation. In the melanoma immunotherapy cohort^35^, cell-level analysis produced stronger apparent statistical evidence than participant-level analysis despite the very small number of independent paired participants, confirming that pseudoreplication can materially distort inference when the experimental unit is ignored. Across the other four datasets, *sctrial* accommodated repeated-measures, paired, cross-sectional, and temporally stratified analyses within a single implementation while preserving design-specific interpretation. Therefore, the principal contribution of *sctrial* is the combination of design-specific estimands, small-sample-aware inference, and a unified workflow that carries study design through estimation, diagnostics, enrichment, and power approximation, which goes beyond participant-aware aggregation alone^17,18^.

Within this framework, our adaptation of difference-in-differences (DiD) to single-cell transcriptomics addresses a specific gap in the existing analytical landscape. DiD has deep roots in econometrics and policy evaluation^29–31^, and its appeal in longitudinal biomedical studies lies in its focus on within-participant change rather than cross-sectional group differences alone. Existing pseudobulk differential expression tools such as muscat^24^ and limma-voom^51,52^ perform inference at the replicate level, and their underlying linear model frameworks can in principle accommodate longitudinal contrasts through interaction terms and subject blocking. In practice, however, these workflows are primarily oriented toward cross-sectional comparisons, and longitudinal applications require manual specification of design matrices, contrasts, and blocking structures that are rarely implemented in published single-cell analyses. Moreover, they return pooled-model coefficients rather than participant-level contrasts, making it difficult to inspect individual trajectories or perform subject-level sensitivity analyses such as leave-one-out diagnostics and permutation tests. *sctrial* formalizes DiD as the default estimand for two-group longitudinal designs and couples it to diagnostics that are natural to a participant-level framing. Nevertheless, the interpretation of DiD depends on substantive assumptions, most notably parallel trends, the assumption that the two groups would have followed similar trajectories in the absence of treatment^53^. This assumption cannot be directly verified with only two time points, though *sctrial* provides a pre-trend interaction test when three or more pre-treatment observations are available. In the analyses presented here, only two time points were available, and baseline group comparisons were therefore used as descriptive diagnostics for baseline comparability. More broadly, *sctrial* reduces pseudoreplication-driven significance inflation and makes the assumptions underlying longitudinal comparisons explicit, enabling investigators to evaluate their plausibility in each study context.

The simulation benchmark situates sctrial’s inferential properties relative to established multi-subject single-cell methods^27,41^. Under pure-null conditions, all evaluated methods were well calibrated and showed comparable power on truly affected genes. The key divergence emerged in mixed-signal panels, where dreamlet^41^ and, to a lesser extent, NEBULA^27^ showed inflated error rates among unaffected genes, a pattern most evident with higher signal fractions and limited numbers of jointly analyzed genes. One possible explanation is that cross-gene borrowing and panel-level normalization become less stable in small gene panels. *sctrial* avoids this dependence through gene-by-gene estimation, at the cost of forgoing the efficiency gains that moderation provides when its assumptions hold. This tradeoff is specific to the evaluated pipeline configurations and should not be generalized to all uses of those methods.

Although constrained in precision by the available sample sizes, the participant-level biological patterns are still informative. In the melanoma cohort, responder-associated trends aligned with B-lineage and humoral programs, whereas non-responder-associated trends aligned with inflammatory and myeloid-associated programs, with cell-type-resolved analysis localizing these signals to B cells/plasma cells and monocytes/macrophages, respectively. The recurring appearance of cytotoxic T cell-associated signals across several intervention-related datasets is consistent with prior work linking T cell functional state to effective anti-tumor immunity^54–56^. The temporal analyses in COVID-19 further show that severity-associated immune differences evolve across the disease course rather than conforming to a single cross-sectional snapshot^57,58^, highlighting the value of time-resolved participant-level analysis.

Several limitations should be noted. The current DiD implementation is centered on binary group comparisons; multi-arm, factorial, or dose-response settings require additional development. Additionally, not all datasets analyzed here represent randomized interventions. In particular, the COVID-19 severity analysis is observational, and severity-associated differences may reflect confounding by clinical or demographic variables in addition to disease biology. In such settings, participant-level estimation remains descriptively useful, but causal interpretation requires assumptions beyond those invoked in a simple longitudinal contrast. Inferential precision also remains constrained by the number of participants. Wild cluster bootstrap procedures and leave-one-out diagnostics help characterize uncertainty but do not overcome limited effective sample size, and our empirical power analyses should therefore be interpreted as data-conditional approximations for study planning. Finally, whole-sample pseudobulk aggregation also means that aggregate effects may reflect both cell-intrinsic transcriptional shifts and changes in cell type composition. While cell type-stratified analyses help address this issue, they do not fully separate state changes from compositional effects.

*sctrial* is released as an open-source Python package integrated with the AnnData/scverse ecosystem to make design-aware participant-level inference practical for repeated-measures single-cell datasets. Extending sctrial to support multi-arm and factorial designs, integrating more tightly with causal-inference workflows for observational studies, and adapting the same principles to spatial transcriptomics would further broaden its scope. Integration with perturbation analysis tools such as pertpy^59^ can also expand its practical applicability. More broadly, this work argues that repeated-measures single-cell studies should be analyzed not only with the correct unit of replication, but with estimands and diagnostics that are explicitly matched to study design

## METHODS

### Datasets

We analyzed five publicly available scRNA-seq datasets spanning melanoma immunotherapy^35^, COVID-19 severity^36^, BNT162b2 vaccination^37^, AML chemotherapy^38^, and CAR-T therapy^39^. For each dataset, the source cohort description is distinguished from the analyzable subset used in this study. A participant was considered analyzable only if the required metadata were available and the participant contributed the samples necessary for the relevant design-specific comparison, such as paired pre/post measurements for longitudinal analyses or clearly defined group labels for cross-sectional contrasts.

#### Melanoma immunotherapy

We used processed scRNA-seq data from Sade-Feldman et al.^35^ (2018), which profiled immune cells from metastatic melanoma tumor biopsies obtained before and during immune checkpoint blockade. The original study analyzed 48 tumor biopsies from 32 patients, including a longitudinal subset with pre-treatment and on-treatment samples. Patients were classified according to clinical response using RECIST-based outcome categories^60^ reported in the source study. For the present work, we restricted the longitudinal DiD analysis to the subset of participants with analyzable paired pre-treatment and on-treatment samples and complete response-group labels. The exact number retained after preprocessing and design-specific filtering is reported in **Table 1**.

#### COVID-19 severity

Single cell PBMC data from Stephenson et al. (2021)^36^ were derived from 143 patients with COVID-19 across a spectrum of disease severities. Severity labels were based on the WHO ordinal scale, and we classified severe disease as WHO score ≥ 5 and mild disease as WHO score ≤ 4. To examine temporal variation in severity-associated immune differences, samples were stratified by days from symptom onset into 0-7, 8-14, and 15+ day windows. Cross sectional severity comparisons were then performed within the relevant temporal strata using the subset of participants meeting the required bin-specific inclusion criteria.

#### BNT162b2 vaccination cohort

We processed a single-cell PBMC dataset from GEO accession GSE171964^37^, corresponding to a BNT162b2 vaccination time-course study, which includes participants sampled at Day 0 and Day 7 and is analyzed as a single-arm paired longitudinal design. Because all participants received vaccination, the primary estimand was the within-participant post-vaccination change rather than a treatment effect relative to an unvaccinated control. Participants were restricted to those with both Day 0 and Day 7 samples available, yielding 6 paired participants for the main analysis.

#### CAR-T therapy

We used longitudinal single-cell data from GSE290722, generated from peripheral blood samples collected from patients with large B-cell lymphoma treated with axicabtagene ciloleucel in the ZUMA-1 setting^39^. The dataset includes paired pre-infusion and post-infusion sampling with accompanying cell type annotations and T-cell receptor profiling. Here, we retained the subset of participants with paired samples (n = 31) and complete metadata required for participant-level longitudinal analysis.

#### AML chemotherapy

A single-cell bone marrow dataset from GSE116256 included samples from 16 AML patients at diagnosis together with matched on-treatment samples, as well as healthy donor controls in the source study^38^. Here, we focused on AML participants with paired diagnosis and post-treatment samples suitable for longitudinal participant-level analysis.

#### Data Preprocessing and Cell Type Annotation

For datasets available as raw FASTQ files, gene-expression reads were processed with Cell Ranger v6.0 against the GRCh38 reference transcriptome. For datasets distributed as processed count matrices, publicly available count matrices and accompanying metadata were imported directly and passed through a harmonized downstream analysis workflow. Quality control was performed at the single-cell level before participant-level aggregation. cells with fewer than 500 detected genes and more than 20% mitochondrial reads were excluded as low quality profiles. Putative doublets were identified using Scrublet^61^. Because doublet-score distributions vary across datasets, the Scrublet threshold was determined using dataset-specific score distributions and applied consistently within each cohort; the final thresholds used for each dataset are reported in **Supplementary Figure 1**. Cells retained after filtering were carried forward for normalization and downstream analysis. For analyses performed on single-cell expression values, normalization was carried out using pooled size factor estimation followed by log transformation.

Cell type annotations were derived from the source datasets when available and harmonized across cohorts using the reported marker-based or study-provided labels. For analyses requiring cell type stratification, only cell types with sufficient numbers of cells and analyzable participant-level observations were retained. Dataset-specific annotation summaries are shown in **Supplementary Figure 2**.

#### Pseudobulk Aggregation

Participant-level inference was performed after collapsing cell-level expression to participant-visit summaries. Specifically, raw counts were summed across all cells within each participant-by-visit group using sparse matrix multiplication, followed by counts-per-million (CPM) normalization and log1p transformation, yielding a participant-level expression matrix with dimensions (participant × visit) by gene. For cell type-stratified analyses, the same procedure was applied separately within each cell type. When the number of cells contributing to a participant-visit summary varied substantially across groups, downstream models incorporated weights based on the contributing cell counts so that summaries derived from larger cell pools contributed proportionately more information. These weights were used as an approximation to account for heterogeneity in the precision of participant-level means rather than as exact inverse variance estimates. Sensitivity analyses comparing alternative aggregation choices, including mean versus median aggregation and different transformation schemes, are shown in **Supplementary Figure 4C–D**. To reduce instability from very sparse genes, genes were retained only if they were detected in at least 20% of participant-visit aggregates within a given analysis set. Detection was defined as nonzero aggregated expression after preprocessing. This filtering was applied before downstream differential analysis, pathway analysis, and cell type-stratified modeling.

Gene signature scores were computed using a z-mean procedure implemented within the sctrial workflow. For each dataset, gene expression values were standardized across cells by subtracting the gene-wise mean and dividing by the gene-wise standard deviation, and the signature score for each cell was defined as the mean standardized expression across all available genes in that signature. Genes with near-zero variance were excluded before scoring to avoid numerical instability. Cell-level signature scores were then aggregated to participant-level summaries by averaging across all cells within each participant-visit group, using the same participant-by-visit grouping structure applied to gene-level analyses but replacing the sum-based CPM procedure with a simple mean, because signature scores are already unitless z-scaled quantities that do not require library-size normalization. When required for visualization or descriptive comparison across signatures, participant-level signature values were standardized after aggregation within the relevant analysis subset. This post-aggregation standardization was used only to place signatures on a comparable display scale and does not change the underlying participant-level design structure used for inference.

### Design-Specific Estimation

For two-group longitudinal analyses, we quantified the primary participant-level contrast using a difference-in-differences (DiD) framework. For a given gene or signature, the DiD estimand is the between-group difference in within-participant pre/post change:

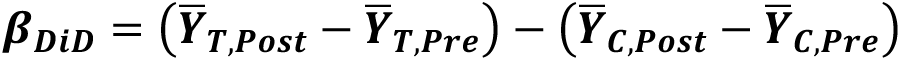

where *T* denotes the focal group, *C* denotes the comparison group, and *Pre* and *Post* denote the two analysis time points. In the melanoma application, these groups corresponded to responders and non-responders, respectively. We use the term DiD effect to emphasize that this quantity represents a group-by-time contrast; its causal interpretation depends on study design and assumptions and is not automatic in observational or response-stratified settings.

In practice, *β_DiD_* was estimated using linear regression with participant fixed effects:

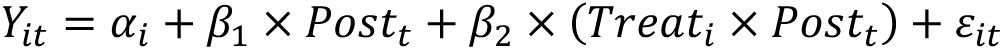

where 𝒀_𝒊𝒕_denotes the participant-level outcome for participant *i* at time point *t* (either aggregated gene expression or a participant-level signature score), 𝛂_𝒊_ is a participant-specific intercept, Post_t_ is an indicator for the post-treatment or follow-up time point, and Treat_i_ × Post_t_ is the group-by-time interaction. Because treatment-group membership is time invariant, its main effect is absorbed by the participant fixed effects and is not separately estimated. The coefficient 𝛃_𝟐_ is therefore the DiD estimand of interest. In the balanced two-period setting, this regression coefficient is algebraically equivalent to the sample DiD contrast above.

Models were fit using ordinary least squares by default. When participant-visit aggregates were based on substantially different numbers of cells, weighted least squares was used with weights proportional to the number of contributing cells as a pragmatic approximation for differing summary precision. SEs were computed using cluster-robust sandwich estimators clustered at the participant level to account for within-participant dependence across repeated measurements. Because cluster-robust inference can be unreliable when the number of clusters is small, we complemented these estimates with wild cluster bootstrap procedures in sensitivity analyses and in the main small sample melanoma analyses.

For single-arm datasets without a concurrent control group (BNT162b2 vaccination, AML chemotherapy, and CAR-T therapy), we quantified participant-level change using paired pre/post contrasts. For each participant and feature, the post-minus-pre difference was computed from participant-level aggregates, and standardized within-arm effect sizes were derived from the distribution of participant-specific differences. For the COVID-19 dataset, which was analyzed as a cross sectional comparison between severity groups within a defined disease time window, we summarized between-group differences using Hedges’ g computed from participant-level values at that window.

Standardized effect sizes were computed to complement the primary participant-level regression estimates. For cross-sectional comparisons, effect size was summarized using Hedges’ g, defined as the standardized difference in group means with small-sample bias correction. For single-arm paired analyses, standardized within-participant change was computed from the distribution of participant-specific post-minus-pre differences. For two-group longitudinal analyses, we additionally summarized descriptive between-group separation by applying standardized mean-difference metrics to the participant-level change scores, while treating the regression-based DiD coefficient as the primary estimand. Confidence intervals for standardized effect sizes were obtained using conventional small-sample procedures appropriate to the corresponding design.

### Bootstrap Inference and Permutation Testing

For DiD analyses in small-sample clustered settings, uncertainty was quantified primarily using a wild cluster bootstrap-t procedure applied at the participant level. For each feature, we first fit the null-imposed model with the DiD interaction constrained to zero, obtained participant-level residuals, and then generated bootstrap samples by multiplying residuals within each participant by independent Rademacher weights (+1 or −1 with equal probability). The perturbed outcome was formed by adding these weighted residuals to the null-imposed fitted values, after which the full DiD model was re-estimated. For each bootstrap replicate, we recorded the studentized statistic obtained by dividing the bootstrap DiD coefficient by its cluster-robust SE. Bootstrap-t confidence intervals were constructed from the empirical quantiles of the bootstrap *t* distribution, and two-sided p-values were computed as the proportion of bootstrap statistics at least as extreme as the observed statistic, using the finite-sample correction (𝑏 + 1)/(𝐵 + 1). We used 1,000 bootstrap iterations unless otherwise stated. Because inference with very small numbers of clusters remains challenging, these procedures were interpreted as small sample-aware approximations rather than exact finite-sample solutions.

To assess whether observed DiD effects exceed what would be expected under random label assignment, we performed participant-level permutation testing in the paired melanoma subset. Response labels were permuted across participants while preserving each participant’s paired pre/post structure, the DiD model was re-estimated for each permutation, and the corresponding test statistic was recorded. The empirical permutation p-value was computed as:

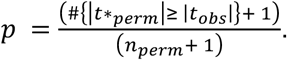

with B = 1,000 permutations.

### Gene Set Enrichment Analysis

For pathway-level analysis, genes were ranked within each cohort using a signed confidence metric defined as 𝑠𝑖𝑔𝑛(𝛽) × (−𝑙𝑜𝑔₁₀(𝑝 + 10⁻¹²)), where β is the design-specific participant-level effect estimate (DiD coefficient for two-group longitudinal analyses, paired participant-level change for single-arm analyses) and p is the corresponding p-value. This metric incorporates both direction and statistical confidence into a single ranking. Pre-ranked gene set enrichment analysis was performed using the GSEApy^43^ package against MSigDB Hallmark^44^, KEGG^45^, Reactome^46^, GO Biological Process^47^, and WikiPathways^48^ collections. Enrichment significance was determined by permutation (1,000 permutations) with FDR correction. For cross-dataset pathway comparison, GSEA was run independently within each cohort, and pathways reaching FDR < 0.25 in at least two datasets were identified as recurrently enriched.

### Robustness and Sensitivity Analyses

To assess sensitivity to individual participants, we performed leave-one-out re-estimation in which each participant was excluded in turn and the participant-level effect was recomputed on the remaining data. Influence was summarized as the change in the estimated effect relative to the full-sample estimate, optionally standardized by the full-sample standard error for comparability across features. We also recorded the proportion of leave-one-out estimates retaining the same sign as the full-sample estimate as a descriptive stability metric.

We also assessed baseline comparability as a descriptive diagnostic for the parallel trends assumption, by comparing pre-treatment participant-level signature values between groups to identify large baseline imbalances that might signal non-comparability.

## Stratified Analysis

For cell-type-specific analyses, cells were first partitioned according to the annotated cell-type labels retained after preprocessing. Participant-level aggregation and the corresponding design-specific analysis were then performed independently within each cell type. Only cell types with sufficient cell counts and analyzable participant-level observations were carried forward. For the resulting cell type × feature tests, p-values were adjusted using the Benjamini–Hochberg procedure over the full set of tests within the relevant analysis.

To examine time-varying severity differences in the COVID-19 severity cohort, samples were stratified by days from symptom onset into early (0-7 days), intermediate (8-14 days), and late (15+ days) bins. Within each bin, we computed participant-level signature summaries separately for severe and mild disease. Severity divergence was defined as the difference in mean participant-level signature score between severe and mild groups within the same bin, whereas Hedges’ g provided a standardized effect size summary for the same contrast. While this binning strategy reduced temporal resolution, it provided interpretable phase-specific summaries of how severity-associated immune differences changed over the disease course.

### Simulation Study and Empirical Power Analysis

To assess Type I error control, statistical power, and calibration under known ground truth, we conducted a Monte Carlo simulation using a hierarchical gamma-Poisson generative model. Gene-level baseline expression rates were drawn from a log-normal distribution calibrated to a real scRNA-seq dataset (melanoma cohort; baseline mean log-rate = −12.86, SD = 2.67), and per-gene overdispersion parameters were drawn from a gamma distribution (median = 0.136). For each participant-visit combination, cell-level UMI counts were generated from a compound gamma-Poisson model, then summed to pseudobulk counts and normalized. Simulations were run across a factorial grid of sample sizes (n = 8, 12, 20, 40, and 60 participants per arm for two-arm designs; equivalent sizes for single-arm paired designs), true DiD effect sizes (β = 0, 0.2, 0.5, and 1.0 on the log-expression scale), and robustness scenarios (heterogeneous dispersion, missing data, cell-count variation, arm imbalance). Each configuration was replicated 200 times with 50 genes per replicate, of which 10 carried the specified effect (20% signal fraction). Four methods were applied to each simulated dataset: (i) *sctrial* DiD: the two-arm *sctrial* estimator was implemented in its algebraically equivalent first-difference form, comparing participant-specific post-minus-pre change scores between arms via Welch’s t-test; (ii) dreamlet^41^: a precision-weighted linear mixed model with voom transformation and empirical Bayes variance moderation; (iii) NEBULA^27^: a negative binomial mixed model operating on cell-level counts with subject-level random effects; and (iv) a nonparametric comparator applied to participant-level change scores (Mann–Whitney U for two-arm; Wilcoxon signed-rank for single-arm). R-based methods (dreamlet, NEBULA) were called via standalone Rscript subprocesses to ensure clean execution environments. For each method and gene, we recorded the estimated effect size, p-value, and runtime. Type I error was defined as the proportion of null genes (true β = 0) with p < 0.05 at each condition. Power was defined as the proportion of signal genes with p < 0.05. Null-gene false positive rate in mixed-signal scenarios was evaluated separately to assess cross-gene contamination from empirical Bayes moderation. P-value calibration was assessed using quantile–quantile plots of observed versus expected −log₁₀(p) for null genes, stratified by scenario type (pure-null versus mixed-signal). Genomic inflation factor (*λ_GC_*) was computed as the ratio of the median observed chi-squared statistic to the expected median under the null.

Two complementary approaches were used to characterize statistical power. First, for the empirical power curves shown in the Results, we repeatedly subsampled participant-level data within each cohort and recomputed the prespecified primary endpoint at a range of sample sizes. For each subsample size, empirical power was defined as the proportion of subsampling iterations yielding nominal significance, and isotonic regression was used to enforce monotonicity of the estimated power curve. Second, for paired and two-group longitudinal settings, the package also provides conventional analytical power and sample-size utilities based on standardized effect sizes, with optional adjustment for clustering through a design-effect factor.

### Software Implementation

*sctrial* is implemented in Python 3.9+ with dependencies on NumPy, Pandas, SciPy, Statsmodels, and AnnData. Optional components support Bayesian DiD estimation via PyMC and mixed-effects modeling through Statsmodels. The package includes convenience functions for rapid analysis together with lower-level workflow interfaces for customized pipelines. Runtime scaling benchmarks are provided in **Supplementary Figure 3J**. Source code, documentation, and reproducible tutorials are publicly available through the package’s GitHub repository: https://github.com/TheOmarLab/sctrial.

## DATA AND CODE AVAILABILITY

Processed single-cell data from Sade-Feldman et al. and Stephenson et al. are available through their original publications. The BNT162b2 vaccination dataset (GSE171964) is available through GEO. Pre-processed analysis-ready datasets are provided through the sctrial package for benchmarking and tutorial purposes. sctrial is available at https://github.com/TheOmarLab/sctrial under the MIT license. Analysis scripts reproducing all figures are provided in the Supplementary Materials.

## ACKNOWLEDGMENTS

We thank the patients and healthy volunteers who contributed samples to the datasets analyzed in this study. We acknowledge the original authors of the public datasets for making their data available.

## AUTHOR CONTRIBUTIONS

M.O. conceived and supervised the project. P.V. and M.O. developed the sctrial software package. I.V. and P.V. performed computational analyses. M.O. and P.V. wrote the manuscript draft. M.R.A. and B.M edited the draft. All authors have read and approved the submitted manuscript.

## SUPPLEMENTARY MATERIAL

### Supplementary Figures

**Supplementary Figure 1. Data quality and cohort characterization. *(A)*** Grouped bar chart showing the number of paired versus unpaired participants in each dataset. ***(B)*** Faceted bar charts showing participant counts for each arm–visit combination. ***(C)*** Box plots of log₁₀(cells per participant + 1) stratified by dataset and arm. ***(D)*** Violin plots of the number of genes detected per cell, stratified by dataset and group. ***(E)*** Violin plots of total counts per cell (left) and percentage of mitochondrial and ribosomal reads (right) with quality control (QC) thresholds. ***(F)*** Lorenz curves and Gini coefficients quantifying the unevenness of cell sampling across participants within each dataset. ***(G)*** Bar chart showing the number of cells retained at each successive QC threshold. ***(H)*** The fraction of participants with cells at each visit across datasets.

**Supplementary Figure 2. Cell type annotation and baseline comparability. *(A)*** UMAP embeddings colored by cell type for each dataset. ***(B)*** UMAP embeddings colored by treatment arm or visit for each dataset. ***(C)*** Dot plot of canonical marker genes for all cell types. For each cell type, the top three marker genes are shown with dot size proportional to the fraction of cells expressing the gene and color intensity proportional to mean z-scored expression. ***(D)*** Bar plots of embedding quality metrics: centroid-silhouette scores (left) and k-nearest-neighbor label purity (right) per dataset. ***(E)*** Heatmap of the fraction of cells assigned to each harmonized cell type within each dataset. ***(F)*** Bar plots showing quantitative arm-mixing score (k = 15, participant-weighted) for each dataset, measuring the degree of treatment-arm separation in the pre-treatment embedding. ***(G)*** Per-dataset parallel categories diagrams showing the flow from participants to cell types, weighted by cell count.

**Supplementary Figure 3. Model diagnostics and assumption checks. *(A)*** Q–Q plots of OLS residuals for each dataset, assessing normality of the residual distribution. ***(B)*** Residual versus fitted value plots across datasets, assessing heteroscedasticity. ***(C)*** Box plots of Cook’s distance analog per dataset, identifying participants with disproportionate influence on parameter estimates. ***(D)*** Scatter plots comparing pre-treatment mean expression between treatment arms (control vs. treated) for each feature in two-arm datasets, and pre-versus post-treatment means for single-arm datasets. Points near the diagonal indicate comparable baseline levels between groups; each feature is labelled. ***(E)*** Signal enrichment quantile–quantile plot. Observed effect quantiles versus permutation null effect quantiles, assessing whether the distribution of effects exceeds chance expectation. ***(F)*** Shapiro–Wilk *W* statistic for normality (left) and Breusch–Pagan statistic for heteroscedasticity (right) summarized across features. ***(G)*** Funnel plot of model effects versus standard errors across features, assessing potential estimation bias. ***(H)*** Lines plots comparing the observed rejection rate versus nominal α from permutation testing, with the identity line representing ideal calibration. ***(I)*** Cleveland dot plot of wall time in seconds across datasets. ***(J)*** Pseudoreplication diagnostics. For each dataset, three-panel comparison showing cell-level versus participant-level β estimates (left), –log₁₀(*p*) values (middle), and standard errors (right).

**Supplementary Figure 4. Sensitivity, robustness, and simulation-based validation. *(A)*** Analytical versus bootstrap standard errors (SEs) across all five datasets. ***(B)*** Standardized versus unstandardized effect sizes in the melanoma cohort. ***(C)*** Scatter plot comparing DiD coefficients from mean versus median pseudobulk aggregation. ***(D)*** Scatter plot comparing DiD coefficients from log₁*p*-transformed versus raw TPM expression. ***(E)*** Heatmap of DiD coefficients across cell types and immune signatures in the melanoma cohort, revealing cell type-specific contributions. ***(F)*** Bar plot of rank-order concordance. Spearman ρ between feature rankings under different preprocessing choices (aggregation method, normalization, standardization) versus the reference pipeline. ***(G)*** Heatmap of maximum leave one out (LOO) deviation across features and datasets. ***(H)*** Two-arm simulation power curves for signal-gene detection (p < 0.05) as a function of true effect size (β), faceted by sample size (n = 20, 40, and 60 per arm), comparing sctrial DiD, dreamlet, NEBULA, and paired Wilcoxon. ***(I)*** Empirical Type I error rates (p < 0.05) for each method under pure-null conditions (β = 0) across sample sizes (n = 8–60 per arm) and both two-arm and single-arm study designs; the dashed red line marks the nominal 5% level. ***(J)*** Genomic inflation factor (*λ_GC_*) under null conditions as a function of sample size for the two-arm design. Values near 1.0 indicate well-calibrated p-value distributions. ***(K)*** Quantile–quantile plots of observed versus expected −log₁₀(p) for null genes, faceted by method. Points are colored by scenario type: method color for pure-null scenarios and gray for null genes in mixed signal (DE) scenarios. ***(L)*** Null-gene false positive rate in mixed signal scenarios as a function of signal-gene effect size (β), at n = 40 per arm (two-arm design). The dashed red line marks the nominal 5% threshold. ***(M)*** Single-arm paired power curves for signal-gene detection as a function of true effect size (*β*), faceted by sample size. ***(N)*** Log-scale box plots of per-iteration wall-clock runtime (including method invocation and I/O) in seconds across all simulation conditions.

**Supplementary Figure 5. Cross-dataset biological consistency. *(A)*** Violin plots of within-dataset z-scored signature scores across all five cohorts. ***(B)*** Heatmap showing pairwise Pearson correlations of standardized effect sizes between all dataset pairs. ***(C)*** Shared top genes with concordant direction. For genes exceeding an effect-size threshold (β > 0.05) in multiple datasets, the frequency of concordant direction is plotted against mean effect. ***(D)*** Violin plots of DiD or Δ effect sizes at the gene level within each cohort. ***(E)*** Forest plot showing the mean T-cell exhaustion effect for the top five cell types within each dataset. For two-arm designs, the effect is the treated-minus-control difference in within-participant change; for single-arm designs, it is the mean within-participant change. Error bars indicate 95% confidence intervals. Points are colored by dataset. ***(F)*** Heatmap of standardized effect sizes for all features and datasets. ***(G)*** Scatter plots of pre-versus post-treatment signature scores with lines connecting paired participants, stratified by dataset. ***(H)*** Heatmap of within-dataset z-scored enrichment results across gene-set libraries, summarizing pathway-level patterns.

**Supplementary Figure 6. Participant-level heterogeneity and temporal dynamics. *(A)*** Raincloud plots of per-participant effects (Post − Pre) in the melanoma cohort for immune-related genes. Each gene shows a half-violin, jittered strip plot, and narrow box plot, stratified by response status (responder in blue, non-responder in orange). The horizontal dashed line indicates zero change. ***(B)*** Heatmap of individual participant effects across immune-related genes. ***(C)*** Response-stratified box plots with Hedges’ *g* annotations, comparing responder and non-responder effect distributions for each gene. ***(D)*** Fraction of total sum of squares attributable to response status (ANOVA η²) for each gene, quantifying the proportion of participant-level variability explained by clinical response. ***(E)*** Fraction of participants with positive effects at each effect size threshold, assessing within-group directional consistency. ***(F)*** Standard deviation (SD) of participant-level effects for each dataset, summarizing inter-individual variability. ***(G)*** Scatter plots of participant-effect SDs between dataset pairs, assessing whether features with high variability in one context also show high variability in another. ***(H)*** Melanoma within-arm change profile. Dot-and-whisker plot of the mean participant-level effect (Post − Pre) ± 95% confidence interval for the top 12 features ranked by absolute effect magnitude, stratified by treatment arm (Responder vs. Non-responder). The dashed line at zero indicates no change. ***(I)*** Single-arm within-arm change profiles. Dot-and-whisker plot of the mean participant-level effect ± 95% confidence interval for the top 12 features across single-arm cohorts (AML, CAR-T, Vaccine), color-coded by dataset. The dashed line at zero indicates no change. ***(J)*** Heatmap of pairwise Pearson correlation coefficients between treated-arm mean fold changes across datasets, with color scale ranging from −1 (blue) to +1 (red) and annotated *r* values.

## Supplementary Tables

**Supplementary Table 1. Gene signature definitions.** Each row corresponds to one immune or clinical gene signature used throughout the study. Columns: Signature (internal identifier), Display Name (human-readable label), Set (Main or Clinical), Gene Count (number of genes in the signature), Genes (comma-separated gene symbols), and Genes_Present_{Dataset} (fraction of signature genes detected in each of the five datasets: Melanoma, COVID-19, Vaccine, AML, CAR-T).

**Supplementary Table 2. Complete effect size results across all datasets.** Unified table of participant-level effect estimates for all immune and clinical signatures across the five cohorts. Columns: dataset, estimand (DiD, within-arm Δ, or Hedges’ *g*), label (display name), feature (signature identifier), effect (standardized coefficient), se (standard error), p-value, FDR (Benjamini–Hochberg), n_units (number of analyzable participants), cov_type_used (covariance estimator), ci_lower, and ci_upper (95% confidence interval bounds). The melanoma cohort uses DiD (responder vs. non-responder); COVID-19 uses cross sectional Hedges’ *g* at DFO 8–14 days; Vaccine, AML, and CAR-T use within-arm paired change.

**Supplementary Table 3. Gene set enrichment analysis (GSEA) results.** Excel workbook with one sheet per dataset (Melanoma, COVID-19, Vaccine, AML, CAR-T). Each sheet reports GSEA pre-ranked results with columns: Name (analysis name), Term (pathway), ES (enrichment score), NES (normalized enrichment score), NOM p-val (nominal p-value), FDR q-val, FWER p-val, Tag % (leading-edge fraction), Gene % (signal fraction), Lead_genes (leading-edge gene list), and Library (gene-set source: Hallmark, Reactome, KEGG, GO BP, WikiPathways). Genes were ranked by the signed confidence metric sign(β) × (−log₁₀(p + 10⁻¹²)).

**Supplementary Table 4. Permutation test results.** Results from 1,000 participant-level label permutations for each signature in each dataset. Columns: dataset, feature, label, observed_beta (observed standardized effect), permutation_p (two-sided empirical p-value), n_permutations (number of valid permutations), null_mean (mean of null distribution), null_sd (standard deviation of null), null_95_lo and null_95_hi (2.5th and 97.5th percentiles of the permutation null), and permutation_FDR (Benjamini–Hochberg corrected within each dataset). For the melanoma cohort, treatment-arm labels were permuted across participants; for single-arm cohorts (Vaccine, AML, CAR-T), visit labels were permuted within participants.

**Supplementary Table 5. Power analysis results.** For each dataset–signature combination, the table reports: observed_effect and observed_se (from Supplementary Table 2), n_participants, sigma_estimated (residual standard deviation estimated from SE and sample size), power_at_observed_N (statistical power at the current sample size), min_N_for_80pct_power (minimum sample size required to achieve 80% power for the observed effect), and min_detectable_effect_80pct (minimum detectable effect at 80% power with the current sample size). Power calculations use the analytical formulas for DiD and paired designs implemented in sctrial.stats.power.

**Supplementary Table 6. Gene-level DiD results for the melanoma cohort.** Genome-wide difference-in-differences (DiD) results for all expressed genes in the Sade-Feldman melanoma dataset. Columns: feature (gene name), beta_DiD (standardized DiD coefficient), se_DiD (standard error), p_DiD (p-value), beta_time (time main effect), p_time, n_units, resid_sd, cov_type_used, FDR_DiD (Benjamini–Hochberg), ci_lower, and ci_upper. Rows are sorted by ascending p_DiD.

