## Supplementary Figures 1-6 for "sctrial: Participant-Level Differential Analysis for Longitudinal Single-Cell Experiments"

### Supplementary Figure 1

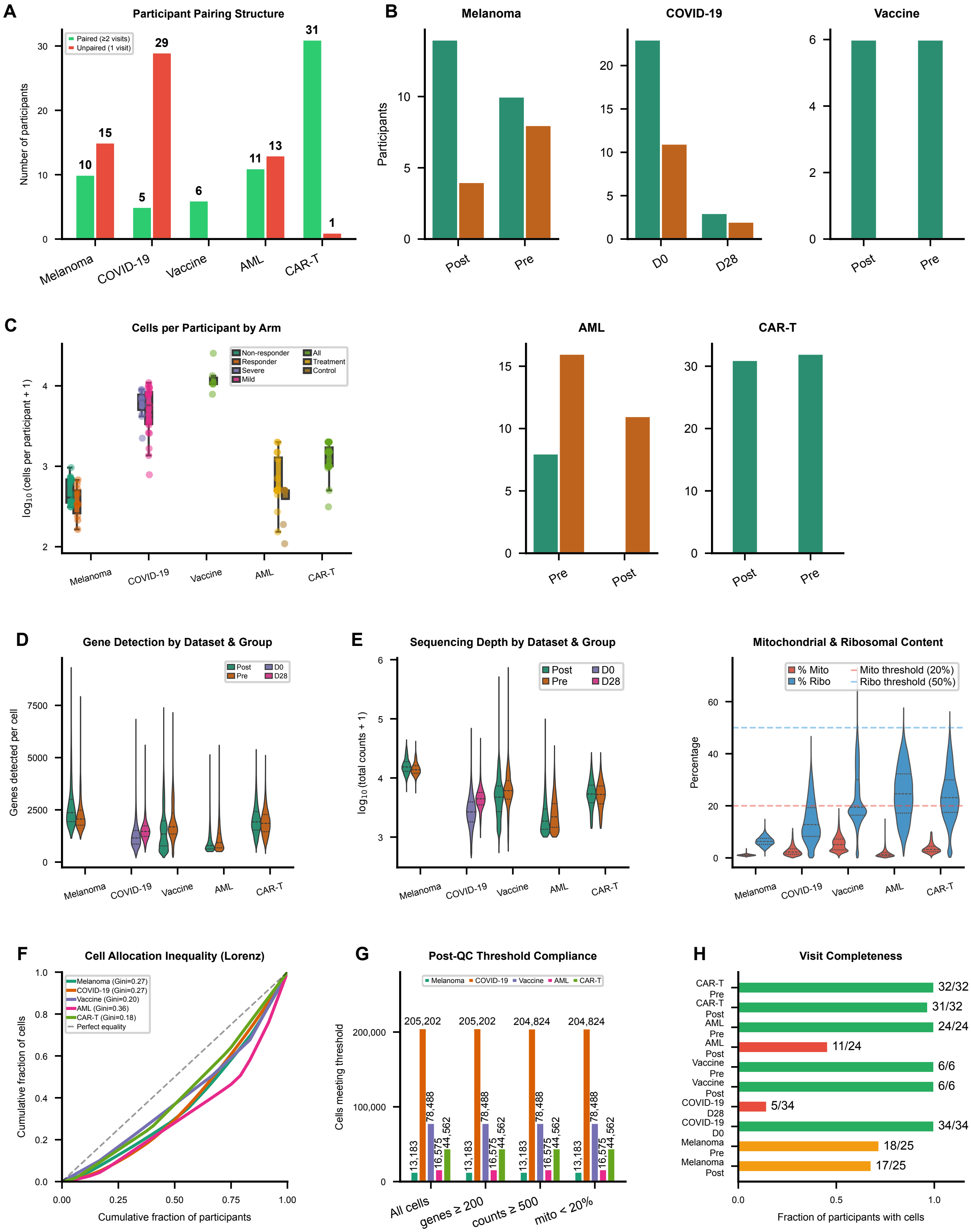

### Supplementary Figure 2

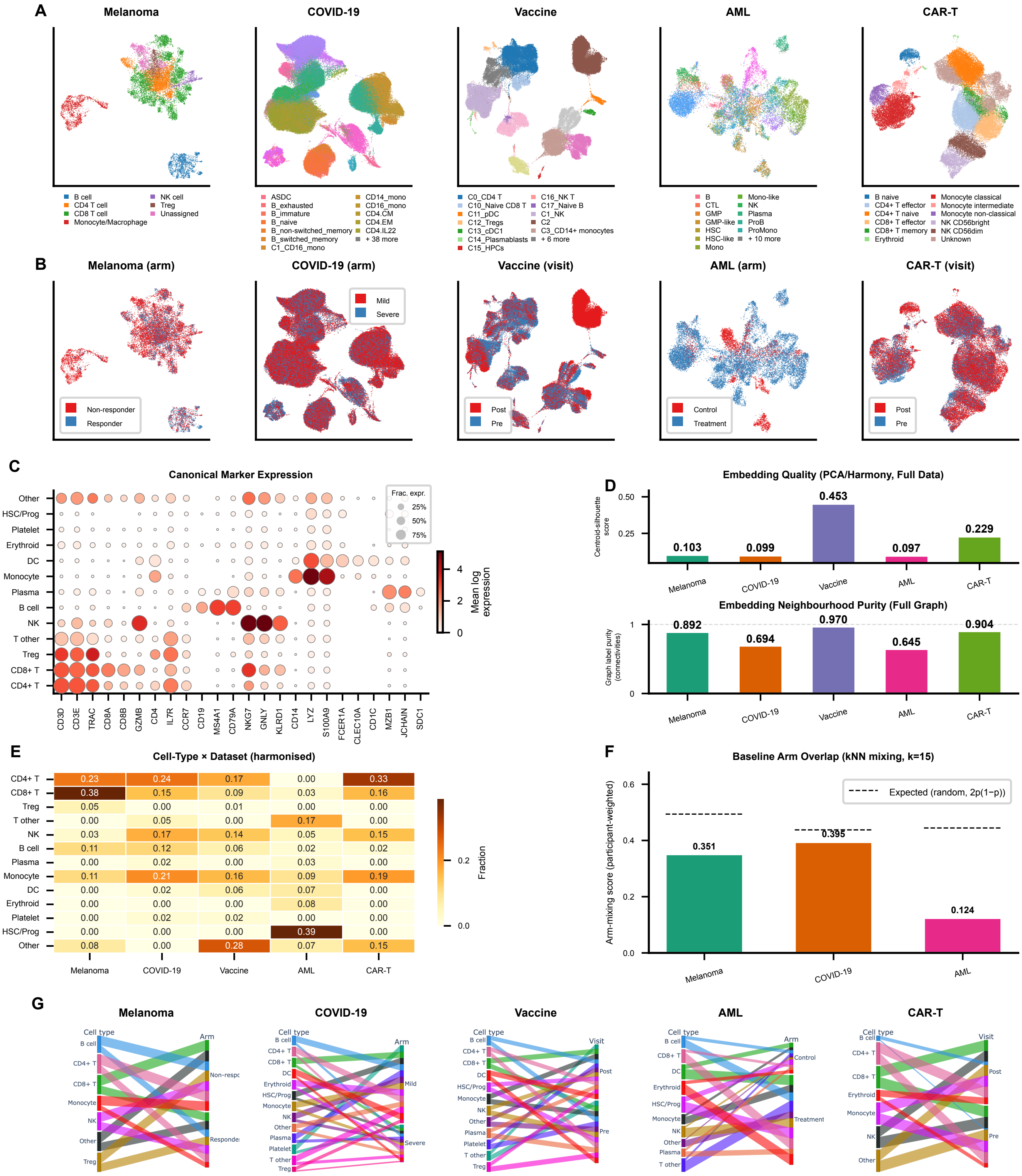


| | Melanoma | COVID-19 | Vaccine | AML | CAR-T |
|---|---|---|---|---|---|
| CD4+ T | 0.23 | 0.24 | 0.17 | 0.00 | 0.33 |
| CD8+ T | 0.38 | 0.15 | 0.09 | 0.03 | 0.16 |
| Treg | 0.05 | 0.00 | 0.01 | 0.00 | 0.00 |
| T other | 0.00 | 0.05 | 0.00 | 0.17 | 0.00 |
| NK | 0.03 | 0.17 | 0.14 | 0.05 | 0.15 |
| B cell | 0.11 | 0.12 | 0.06 | 0.02 | 0.02 |
| Plasma | 0.00 | 0.02 | 0.00 | 0.03 | 0.00 |
| Monocyte | 0.11 | 0.21 | 0.16 | 0.09 | 0.19 |
| DC | 0.00 | 0.02 | 0.06 | 0.07 | 0.00 |
| Erythroid | 0.00 | 0.00 | 0.00 | 0.08 | 0.00 |
| Platelet | 0.00 | 0.02 | 0.02 | 0.00 | 0.00 |
| HSC/Prog | 0.00 | 0.00 | 0.00 | 0.39 | 0.00 |
| Other | 0.08 | 0.00 | 0.28 | 0.07 | 0.15 |

### Supplementary Figure 3

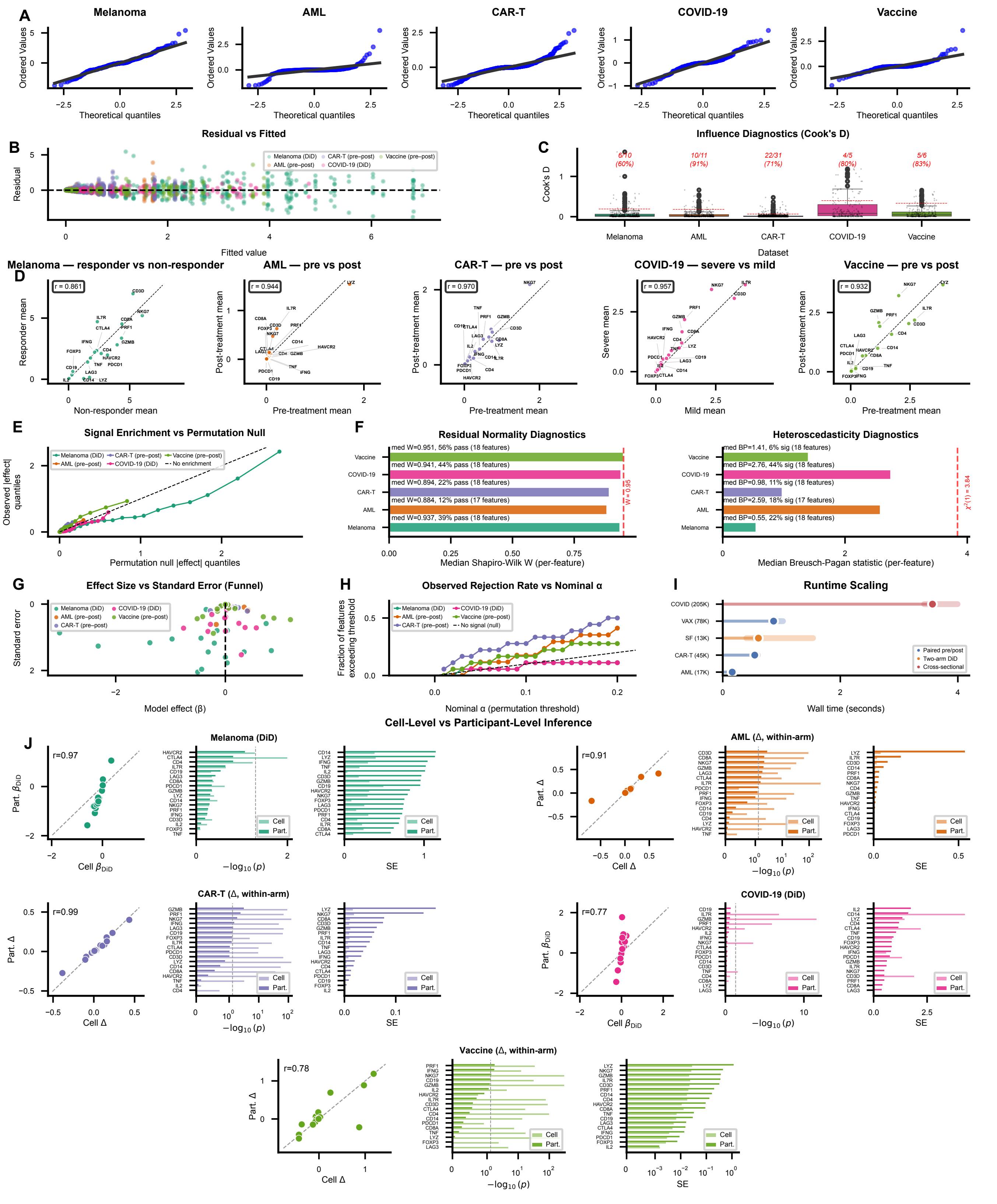

### Supplementary Figure 4

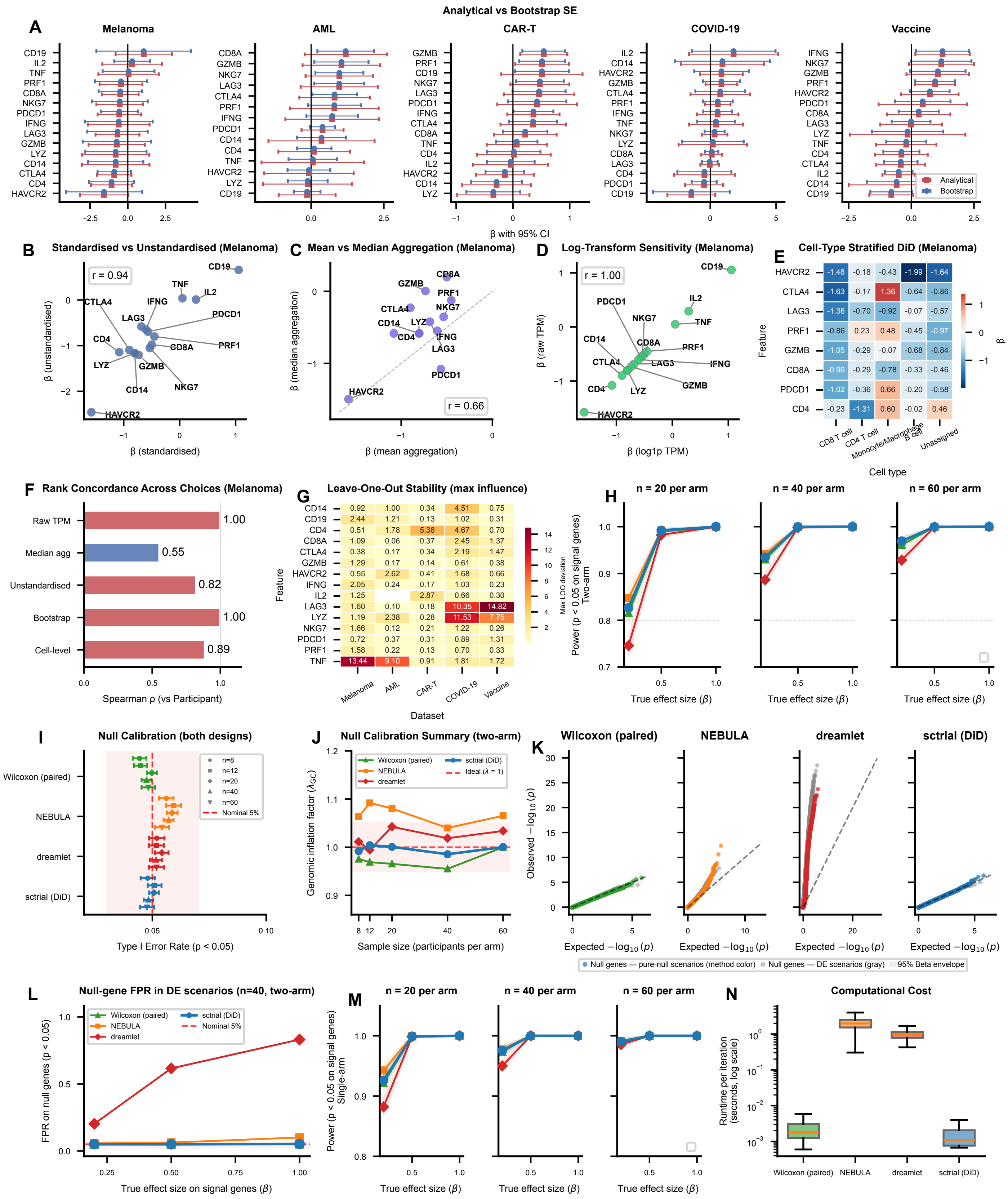

### Supplementary Figure 5

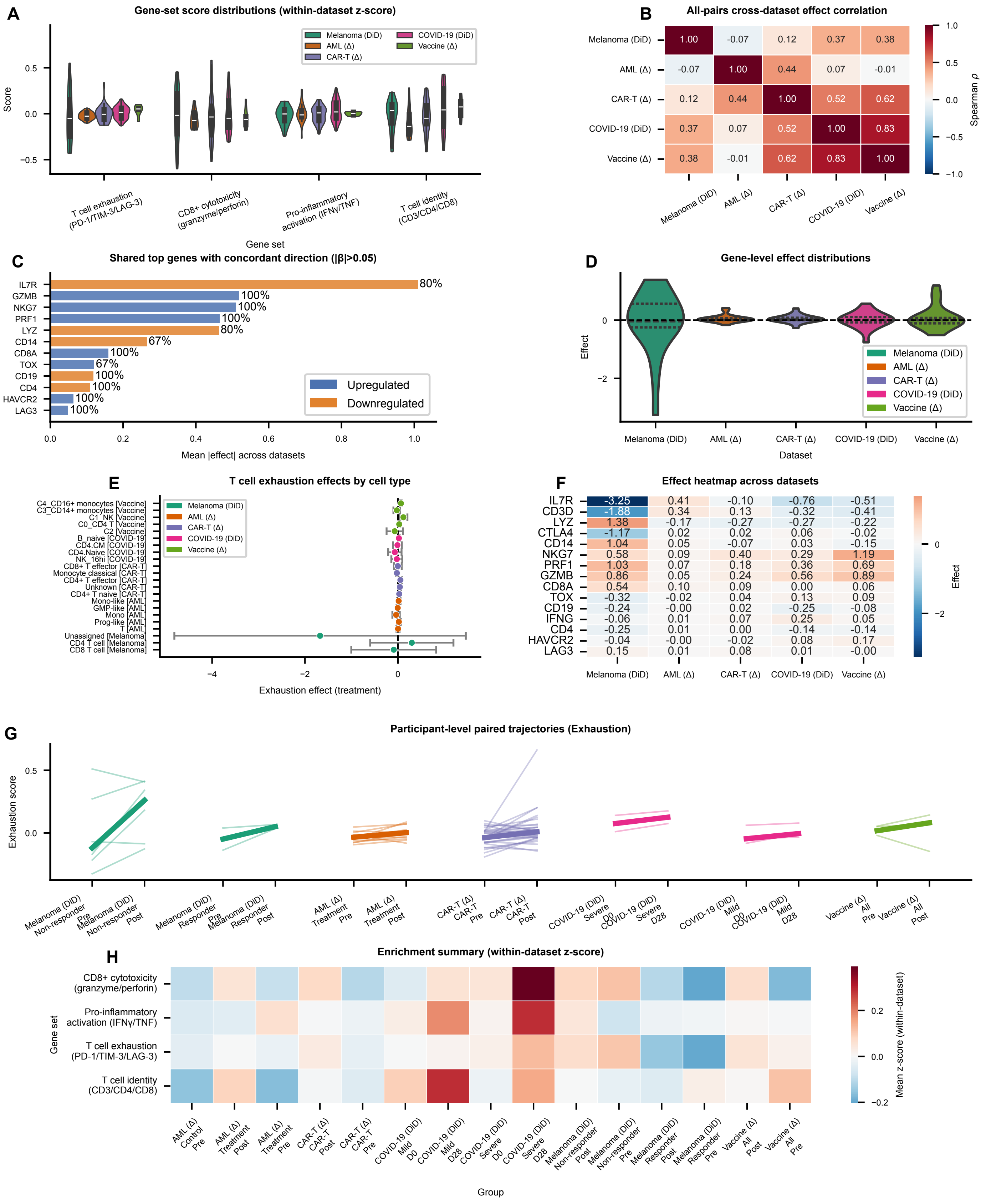

### Supplementary Figure 6

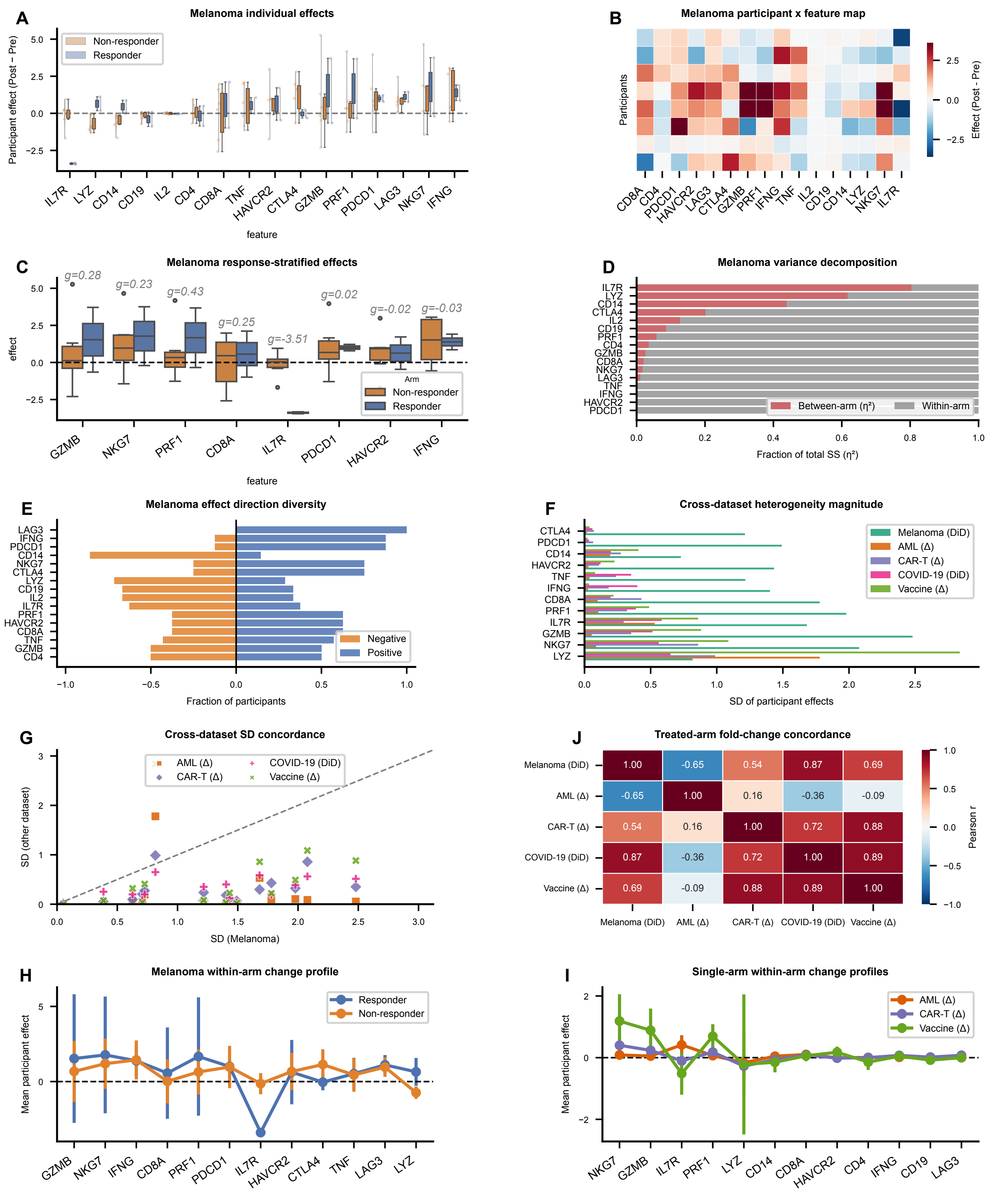
